# Cargo receptor-assisted endoplasmic reticulum export of pathogenic α1-antitrypsin polymers

**DOI:** 10.1101/2020.09.17.301119

**Authors:** Adriana Ordonez, Heather P Harding, Stefan J Marciniak, David Ron

## Abstract

Circulating polymers of alpha1-antitrypsin (α1AT) are chemo-attractant for neutrophils and contribute to inflammation in pulmonary, vascular and adipose tissues. Cellular factors affecting the intracellular itinerary of mutant polymerogenic α1AT remain obscure. Here, we report on an unbiased genome-wide CRISPR/Cas9 screen for regulators of trafficking of the polymerogenic α1AT^H334D^ variant. Single guide RNAs targeting genes whose inactivation enhanced accumulation of polymeric α1AT were enriched by iterative construction of CRISPR libraries based on genomic DNA from fixed cells selected for high polymer content by fluorescence-activated cell sorting. This approach bypassed the limitation to conventional enrichment schemes imposed by cell fixation and identified 121 genes involved in polymer retention at false discovery rate < 0.1. From that set of genes, the pathway ‘cargo loading into COPII-coated vesicles’ was overrepresented with 16 significant genes, including two transmembrane cargo receptors, LMAN1 (ERGIG-53) and SURF4. *LMAN1* and *SURF4*-disrupted cells displayed a secretion defect extended beyond α1AT monomers to polymers, whose low-level secretion was especially dependent on SURF4 and correlated with SURF4-α1AT^H334D^ physical interaction and with enhanced co-localisation of polymeric α1AT^H334D^ with the endoplasmic reticulum (ER). These findings suggest that ER cargo receptors co-ordinate intracellular progression of α1AT out of the ER and modulate the accumulation of polymeric α1AT not only by controlling the concentration of precursor monomers but also through a previously-unrecognised role in secretion of the polymers themselves.

## Introduction

Alpha1-Antitrypsin (α1AT) (*SERPINA1*) is a glycoprotein synthesised primarily in hepatocytes and secreted as a monomer into blood to constitute the most abundant SERine Protease INhibitor (SERPIN) in circulation. Its main function is to inhibit neutrophil elastase in lungs defending against excessive tissue degradation by the endogenous protease-enzyme activity (Carrell and Lomas, 2002).

Missense variants in *SERPINA1*, including the most-common Z variant (E342K), perturb the stability and conformation of α1AT monomers, resulting in their intracellular retention and formation of ordered and pathogenic polymers that accumulate within the lumen of the endoplasmic reticulum (ER) of hepatocytes. Intracellular retention is the basis of plasma α1AT deficiency underlying early-onset emphysema (Gooptu et al., 2014). Accumulation of polymers within liver cells is also associated with a toxic gain-of-function that predisposes to neonatal hepatitis and hepatocellular carcinoma (Eriksson et al., 1986). Interestingly, only 10-15% of patients develop severe liver pathology, suggesting variation in the handling of intracellular polymers (Wu et al., 1994).

Whilst α1AT polymers are most abundant intracellularly, polymers have also been identified in circulation (Tan et al., 2014) and in tissues; in the skin and kidney of α1AT-deficient patients with panniculitis (Gross et al., 2009) or vasculitis (Morris et al., 2011) and in bronchoalveolar lavage fluid of patients with lung disease (Morrison et al., 1987). *In vitro* (Mulgrew et al., 2004) and *in vivo* studies (Mahadeva et al., 2005) implicate extracellular polymers as chemo-attractants for human neutrophils that could contribute to inflammation and lung damage and less common extra-pulmonary manifestations of α1AT deficiency (Gooptu and Lomas, 2008).

Despite its importance to disease development, the processing and fate of intracellular polymers remain poorly understood. Both autophagy and ER-associated degradation (ERAD) have been implicated in their clearance (Kroeger et al., 2009). Less is known about how polymers reach the extracellular compartment. This has long been thought to be the result of either polymer release from dying cells or polymerisation of mutant α1AT secreted as monomers. Recently, studies of plasma of α1AT-deficient patients before and after liver transplant (Tan et al., 2014) and cellular models suggest that circulating polymers are more likely to arise from secretion of pre-formed polymers rather than polymerisation extracellularly (Fra et al., 2016). Notably, levels of polymers in plasma from α1AT-deficient patients do not increase after incubation at 37°C for 3 days (Fra et al., 2016). This observation suggests that plasma levels of mutant polymerogenic α1AT (which are typically 10-15% the levels found in normal individuals) are below the threshold for aggregation. However, the processes underlying polymer secretion remain largely unknown.

Here, we performed a forward genetic screen to identify components affecting the intracellular levels of a highly polymerogenic α1AT variant, the King’s mutant (H334D) (Miranda et al., 2010). Our observations indicate that α1AT polymers can be secreted from the cells by the canonical secretory pathway and identify LMAN1 and SURF4 as cargo receptors involved in the trafficking of monomeric and polymeric α1AT.

## Results

### Flow cytometry-based assay to monitor intracellular α1AT polymers

To identify genes that modify intracellular levels of α1AT polymers, we developed a quantitative fluorescence-activated cell sorting (FACS)-compatible readout for the abundance of intracellular polymers using the well-described α1AT polymer-specific monoclonal antibody 2C1 (Mab2C1) (Miranda et al., 2010) in a previously-characterised CHO-K1 cell line (Ordonez et al., 2013). These cells express the polymerogenic variant (H334D) of α1AT, under control of a tetracycline**-**inducible (Tet-on) promoter that enables tight regulation of α1AT expression (Fig. S1A). A derivative CHO-K1 Tet-on_α1AT^H334D^ clone that stably expresses Cas9 and maintained parental regulation of Tet-inducible α1AT^H334D^ expression was selected for screening.

To favour an experimental system that could respond to genetic perturbations with an increase in intracellular α1AT^H334D^ polymers, cells were treated with a range of concentrations of doxycycline in the absence or presence of BafilomycinA1, an inhibitor of lysosomal activity. BafilomycinA1 enhances accumulation of α1AT polymers (Kroeger et al., 2009) and proved useful in exploring the dynamic range of the assay. Doxycycline at 5-50 ng/ml was associated with low basal levels of Mab2C1-staining that increased conspicuously upon BafilomycinA1 treatment, suggesting a suitable assay window for the screen (Fig. S1B).

### A genome-wide screen identifies a set of genes affecting the intracellular itinerary of polymerogenic α1AT

CHO-K1 Tet-on_α1AT^H334D^_Cas9 cells were initially transduced with a genome-wide CRISPR/Cas9 knockout library (Lib_0_) comprising 125,030 single guide RNAs (sgRNAs) (Fig. 1A). α1AT^H334D^ expression was then induced with doxycycline followed, 24 hrs later by fixation, permeabilisation and staining with the Mab2C1 primary antibody. Cells were FACS sorted into 3 bins based on Mab2C1-dependent fluorescence intensity: ‘brightest’, ‘medium-bright’ and ‘dull’ (Fig. 1B).

**Fig. 1.**
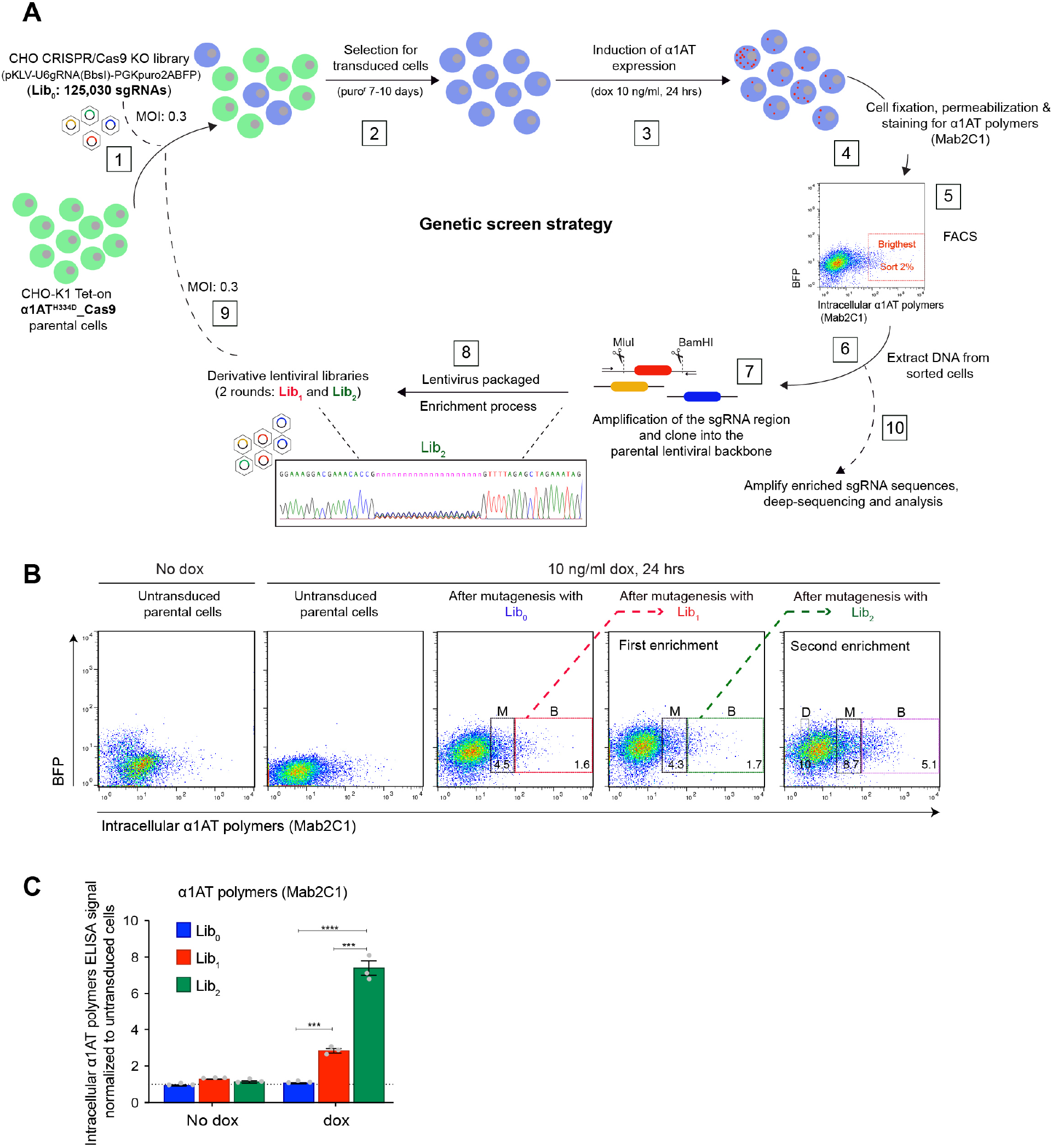
CRISPR/Cas9 screen to identify modifiers of intracellular levels of α1-antitrypsin polymers. **(A)** Workflow of a genome-wide CRISPR/Cas9 knockout (KO) screen. CHO-K1 cells expressing Cas9 and a Tet-inducible allele of α1AT^H334D^ were transduced at low multiplicity of infection (MOI: 0.3) with a lentiviral library of sgRNAs targeting the whole CHO genome (Lib_0_) [1]. Transduced cells were selected for presence of the puromycin resistance marker [2]. Expression of the α1AT^H334D^ transgene was induced with doxycycline (dox) [3]. Cells were fixed and stained for polymeric α1AT using the polymer-specific monoclonal antibody 2C1 (Mab2C1) [4] and FACS sorted based on signal intensity [5]. Genomic DNA was extracted from pools of cells with the highest level of polymer signal (‘brightest’) [6] and used to amplify enriched sgRNA sequences to create new lentiviral libraries (Lib_1_ and Lib_2_). Sanger sequencing indicates the presence of sgRNA sequence diversity in the new lentiviral Lib_2_ [7 & 8]. The selection cycle was repeated [9] and at its conclusion [10] genomic DNA from the selected cells was prepared for high throughput sequencing and analysis of the successively enriched sgRNA sequences. **(B)** Dual-channel flow cytometry of intracellular levels of α1AT polymers (stained with Mab2C1) and BFP (transduction marker) in α1AT^H334D^-expressing cells before and after transduction with Lib_0_ (unenriched library), and successively-enriched Lib_1_ and Lib_2_. The boxed areas include the cells sorted for genomic analysis: ‘brightest’ (B), ‘medium-bright’ (M) and ‘dull’ (D). **(C)** Intracellular α1AT polymer signals quantified by sandwich ELISA of unsorted cells, transduced with Lib_0_, Lib_1_ and Lib_2_, respectively, in the presence or absence of doxycycline (dox; 10 ng/ml, 24 hrs). Shown is the mean ± SEM normalised to untransduced cells of three independent experiments. ***, P < 0.001; and ****, P < 0.0001. Unpaired t-test.

Cell fixation, required to detect intracellular polymers, precluded conventional enrichment schemes through successive rounds of phenotypic selection and expansion of the pooled cells. To circumvent this impasse, we implemented an approach based on recovery of sgRNA sequences from phenotypically-selected cell populations (Fig. 1A,B). Genomic DNA from the ‘brightest’-sorted cells was extracted and fragments covering integrated sgRNA sequences were PCR-amplified and used to generate a derivative CRISPR library (Fig. 1A, lower segment). The derivative library (Lib_1_), enriched in viral particles bearing phenotype-linked sgRNA sequences, was transduced into parental CHO-K1 Tet-on_α1AT^H334D^_Cas9 cells followed by further phenotypic selection and generation of a second, enriched derivative library (Lib_2_, Fig. 1B). Transduction with Lib_0_, Lib_1_ and Lib_2_ progressively increased intracellular α1AT polymers, as assessed by FACS (Fig. 1B) and ELISA (Fig. 1C).

Next, genomic DNA, pooled from sorted cells in the different bins at different stages of the phenotypic enrichment process and from unsorted control cells, was subjected to high-throughput sequencing and MAGeCK bioinformatics analysis (Li et al., 2014) to determine sgRNA sequence enrichment and the corresponding gene ranking list (Table S1). Quality control based on sgRNA sequence read counts showed that over 90% of the reads mapped to the libraries (Fig. S2A). Distribution of normalised read counts indicated that after successive rounds of positive phenotypic selection the diversity of sgRNA species declined from libraries Lib_0_ to Lib_2_, with increasing percentage of sgRNAs with zero read counts and sgRNA with very high counts (Fig. S2B,C).

Gene ontology (GO) analysis of the most significantly enriched genes in the ‘brightest’ Mab2C1-stained cells [with a false discovery rate (FDR) < 0.1] revealed that ‘regulation of chromosome organisation’ was the strongest selected GO term (Fig. 2A). This cluster, thought to reflect the indirect effects of altered transcriptional regulation on polymer levels, was not further considered. The second highly represented cluster was ‘cargo loading into COPII-coated vesicle’, which included 16 genes that were significantly enriched during the selection process (Fig. 2A,B). These encode components of the coat protein II (COPII) complex that initiates vesicle budding at the ER (*SEC23B, SAR1A* and *SEC24B*), non-COPII proteins important to vesicle formation (*RAB1A, TFG, TRAPPC12* and *MAPK10*) (D’Arcangelo et al., 2013) and two cargo receptors with a known role in protein transport from ER to Golgi apparatus (*LMAN1* and *SURF4*) (Gomez-Navarro and Miller, 2016). In addition, protein-protein interaction network analysis of the proteins encoded by the 16 identified genes revealed that 5 of them form an independent network, highlighting their interconnectivity (Fig. 2C). Thus, this screen hints at an important role for the early secretory pathway in specifying intracellular levels α1AT polymers (Fig. 2D).

**Fig. 2.**
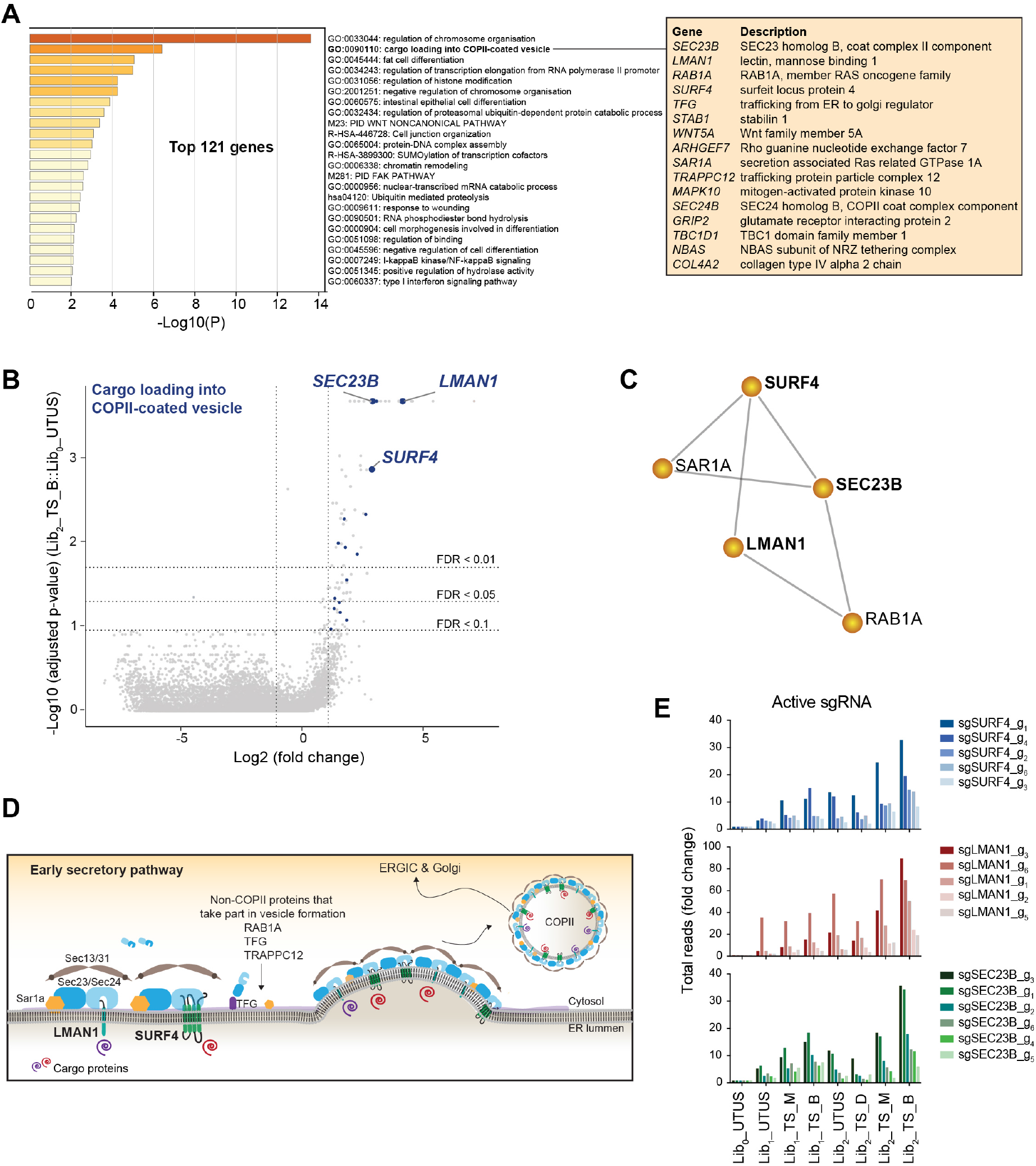
sgRNAs targeting genes encoding components of the early secretory pathway are enriched in cells with elevated intracellular α1-antitrypsin polymers. **(A)** Gene ontology (GO) enrichment analysis of the top 121 hits identified in the CRISPR screen and annotation of the 16 genes included in the GO term ‘cargo loading into COPII-coated vesicle’. **(B)** Volcano plot showing the Log_2_ (fold change) and the Log_10_ (adjusted p-value) of the genes targeted by sgRNAs in ‘treated and sorted’ (TS) cells transduced with Lib_2_ versus ‘untreated and unsorted’ (UTUS) cells transduced with Lib_0_. Genes above the horizontal dashed lines were significantly enriched in Lib_2_. Genes of the GO term ‘cargo loading into COPII-coated vesicle’ are in blue. **(C)** Protein-protein interaction network (Metascape) of the 16 proteins encoded by the genes of the ‘cargo loading into COPII-coated vesicle’ cluster. **(D)** Cartoon of the early secretory pathway where relevant factors identified in the screen are depicted. **(E)** Total reads for each active sgRNA targeting the selected genes for validation.

### Elevated intracellular α1AT^H334D^ polymer levels in cells lacking SURF4, LMAN1 and SEC23B

Of the genes targeted by guides enriched in the ‘brightest’ cells, we deemed those encoding proteins with an ER luminal domain that could interact with polymers to be of particular interest. LMAN1 (lectin mannose binding1) and SURF4 (surfeit protein locus 4), two transmembrane cargo receptors (Hauri et al., 2000; Reeves and Fried, 1995), satisfied that criterion. Another highly enriched gene, *SEC23B*, encoding the cytosolic component of the COPII machinery (Jensen and Schekman, 2011), was included as a reference (Fig. 2A,B). Five of the six sgRNA targeting each of these three genes were significantly enriched in the ‘brightest’ population, adding confidence that they represent reliable hits (Fig. 2E).

To validate the genotype-phenotype relationship suggested by the screen, *SURF4, LMAN1* and *SEC23B* were re-targeted by CRISPR/Cas9-mediated gene disruption in parental CHO-K1 Tet-on_α1AT^H334D^ cells, using two guides mapping to separate exons (Fig. 3A). Cells expressing wild-type α1AT (Ordonez et al., 2013) were also targeted. Clonal knockout derivative cell lines were validated by genomic sequencing and, in case of *SURF4* and *LMAN1*, by evidence for depletion of the proteins by immunoblotting (Fig. 3B,C).

**Fig. 3.**
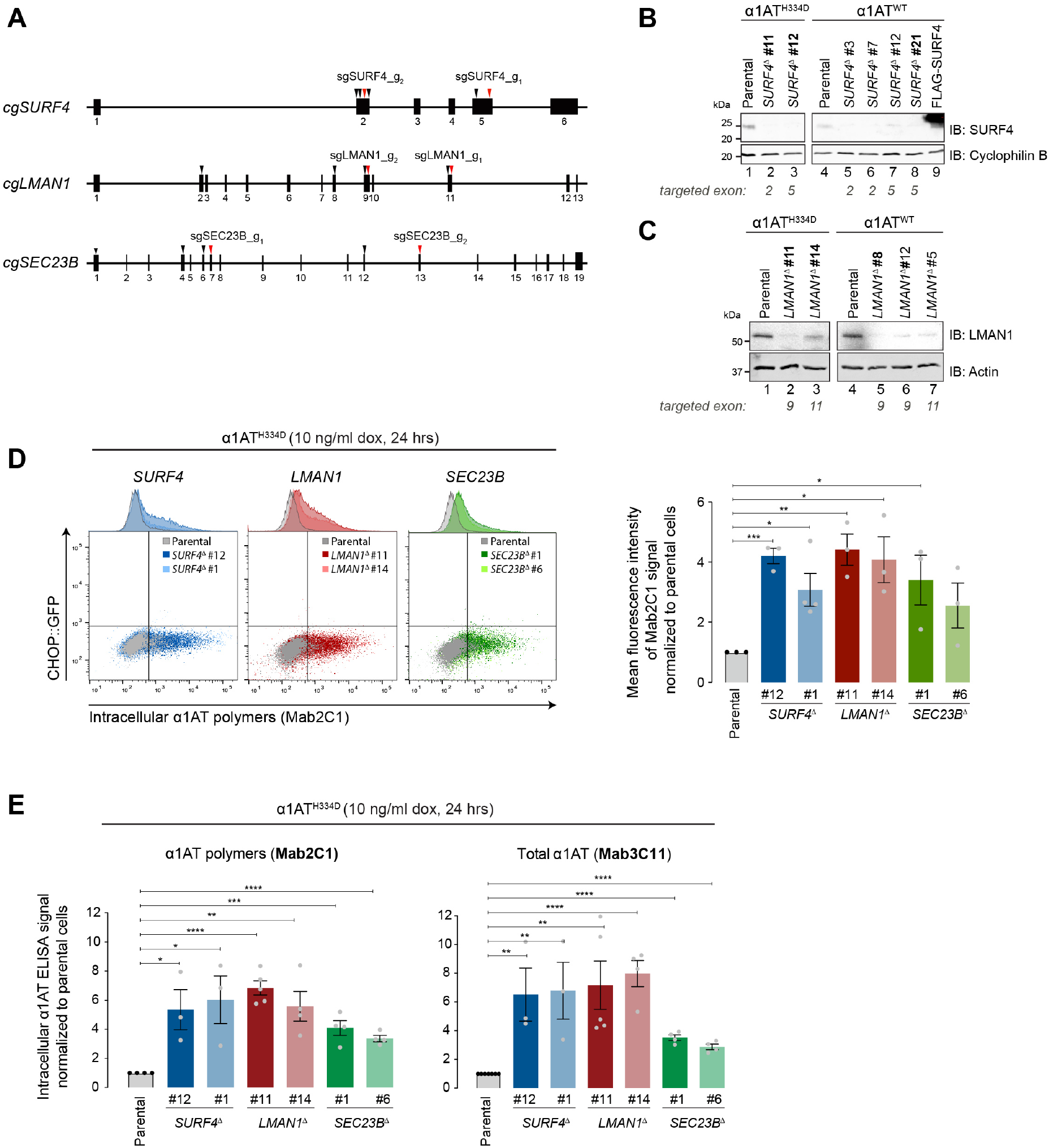
Disruption of *SURF4, LMAN1* and *SEC23B* increases the intracellular levels of α1-antitrypsin polymers in CHO-K1 cells. **(A)** Diagrams of the *Cricetulus griseus SURF4, LMAN1* and *SEC23B* loci showing the target sites of the 6 sgRNAs (arrowheads) included in the CRISPR/Cas9 library. Red arrowheads indicate sgRNAs selected for validation. **(B and C)** Immunoblots of SURF4 (upper panel) and LMAN1 (lower panel) in lysates of parental CHO-K1 Tet-on cells expressing either α1AT^H334D^ or α1AT^WT^ and several *SURF4* and *LMAN1* deleted derivatives. In bold, clones selected for functional experiments. Lysate of parental cells transfected with a FLAG-SURF4-ecoding plasmid served as a positive control. **(D)** Dual-channel flow cytometry of intracellular levels of α1AT polymers and *CHOP::GFP* in CHO-K1 parental Tet-on_α1AT^H334D^ cells and two independent clones where *SURF4, LMAN1* or *SEC23B* were disrupted. The bar graph shows the mean ± SEM of the Mab2C1-signal normalised to doxycycline-treated parental cells from three or four independent experiments. **(E)** As in ‘D’ but plotting the intracellular α1AT signal from sandwich ELISA assays using the anti-polymer Mab2C1 (left panel) and the anti-total α1AT Mab3C11 (right panel). *, P < 0.05; **, P < 0.01; ***, P < 0.001; and ****, P < 0.0001. Unpaired t-test.

Disruption of *SURF4, LMAN1* and *SEC23B* increased intracellular polymer levels as assessed by flow cytometry after immunostaining of polymeric α1AT^H334D^ (Fig. 3D). These observations were confirmed by ELISA with two different antibodies: the polymer-specific Mab2C1 and a monoclonal antibody that recognises all α1AT conformers (Mab3C11) (Fig. 3E).

*SURF4* and *LMAN1*, confirmed above as genes whose inactivation enhances levels of intracellular polymeric α1AT^H334D^, play a broad role in trafficking of cargo out of the ER. Perturbations in ER function caused by protein misfolding or by impeded egress of proteins from the ER lead to ER stress and trigger the unfolded protein response (UPR), a protective and adaptive response aimed to re-establish ER homeostasis (Walter and Ron, 2011). Notably, *in vitro* studies indicate that Brefeldin A, an inhibitor of protein transport from ER to the Golgi apparatus, leads to the activation of the UPR (Citterio et al., 2008). Therefore, to gauge the contribution of any general perturbation to ER function that may arise from the inactivation of such genes, we turned to CHO-K1 S21 cells bearing *CHOP::GFP* and *XBP1s::Turquoise* unfolded protein response (UPR) reporters (Sekine et al., 2016). *SURF4* and *LMAN1* were inactivated by sgRNA whose expression was linked to a mCherry reporter. This enabled scoring UPR activation in populations of mutant cells, free of the bias that might otherwise be introduced by clonal selection. No induction of the UPR reporters was observed following *SURF4* and *LMAN1* inactivation. Inactivation of *HSPA5*, encoding the ER chaperone BiP, a positive control, strongly inducing both UPR branches (Fig. 4). These observations indicate that inactivation of *SURF4* and *LMAN1* did not globally perturb ER protein homeostasis and suggested that the observed increase in polymers may arise by a more specific mechanism.

**Fig. 4.**
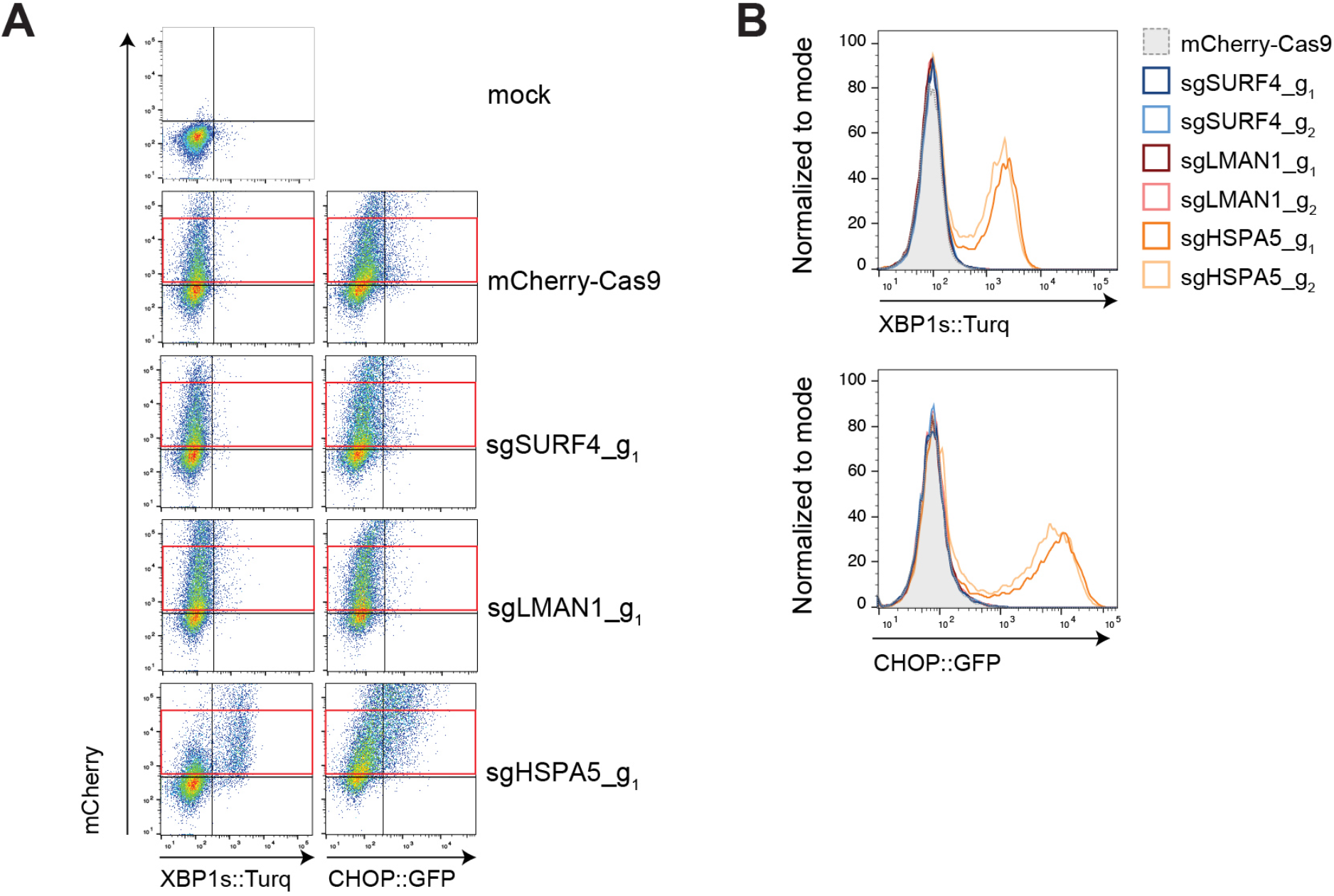
SURF4 and LMAN1 depletion does not activate the unfolded protein response. **(A)** Dual-channel flow cytometry of *XBP1s::Turquoise* or *CHOP::GFP* and mCherry in CHO-K1 S21 cells transiently transfected with sgRNA-mCherry-Cas9 plasmids targeting *SURF4, LMAN1* and *HSPA5* (BiP protein). Dot-plots are representative of one experiment. The red rectangles delineate cells expressing moderate levels of mCherry-tagged plasmid selected for the histogram shown in ‘B’. **(B)** Distribution of the *XBP1s::Turquoise* and *CHOP::GFP* signals, in mCherry-positive cells gated by red rectangles in ‘A’. The same experiment was repeated with equal results using a second sgRNA for each gene.

### LMAN1 and SURF4 promote trafficking of α1AT in CHO-K1 cells

LMAN1 has been previously implicated in mediating ER exit of wild-type monomeric α1AT (Nyfeler et al., 2008; Zhang et al., 2011). SURF4, by contrast, has been reported to lack such a function; at least in HEK293 cells (Emmer et al., 2018). To examine the roles of SURF4 and LMAN1 in the trafficking of polymerogenic α1AT^H334D^, we performed pulse-chase experiments to compare the kinetics of α1AT secretion and the accumulation of polymers in parental, *SURF4*^Δ^ and *LMAN1*^Δ^ CHO-K1 Tet-on_α1AT^H334D^ cells. Cells were pre-treated with low concentration of doxycycline followed by radioactive pulse labelling for 20 min and a subsequent chase (Fig. 5A). α1AT immunoprecipitation from cell-lysates and culture media was performed with antibodies reactive with all forms of α1AT (total) or selective for polymers (Mab2C1) (Fig. 5B). α1AT contains three N-glycosylation sites. Thus, the ER-associated 52-kDa α1AT^H334D^ species gradually appeared in the culture media as mature-glycosylated species of 55-kDa (Fig. 5B). Disruption of *LMAN1*, and to a lesser degree *SURF4*, led to a significant defect in the clearance of the ER form and appearance of the mature-glycosylated form in the culture media (Fig. 5B,C). This trend was even more conspicuous in terms of α1AT^H334D^ polymer secretion as *LMAN1*^Δ^ and *SURF4*^Δ^ cells accumulated more intracellular polymers than parental cells (Fig. 5B,D). Similar findings were observed in an independently derived *SURF4*^Δ^ clone (Fig. S3). Interestingly, both *LMAN1*^Δ^ and *SURF4*^Δ^ cells secreted proportionally fewer α1AT^H334D^ polymers than parental cells (Fig. 5B,E).

**Fig. 5.**
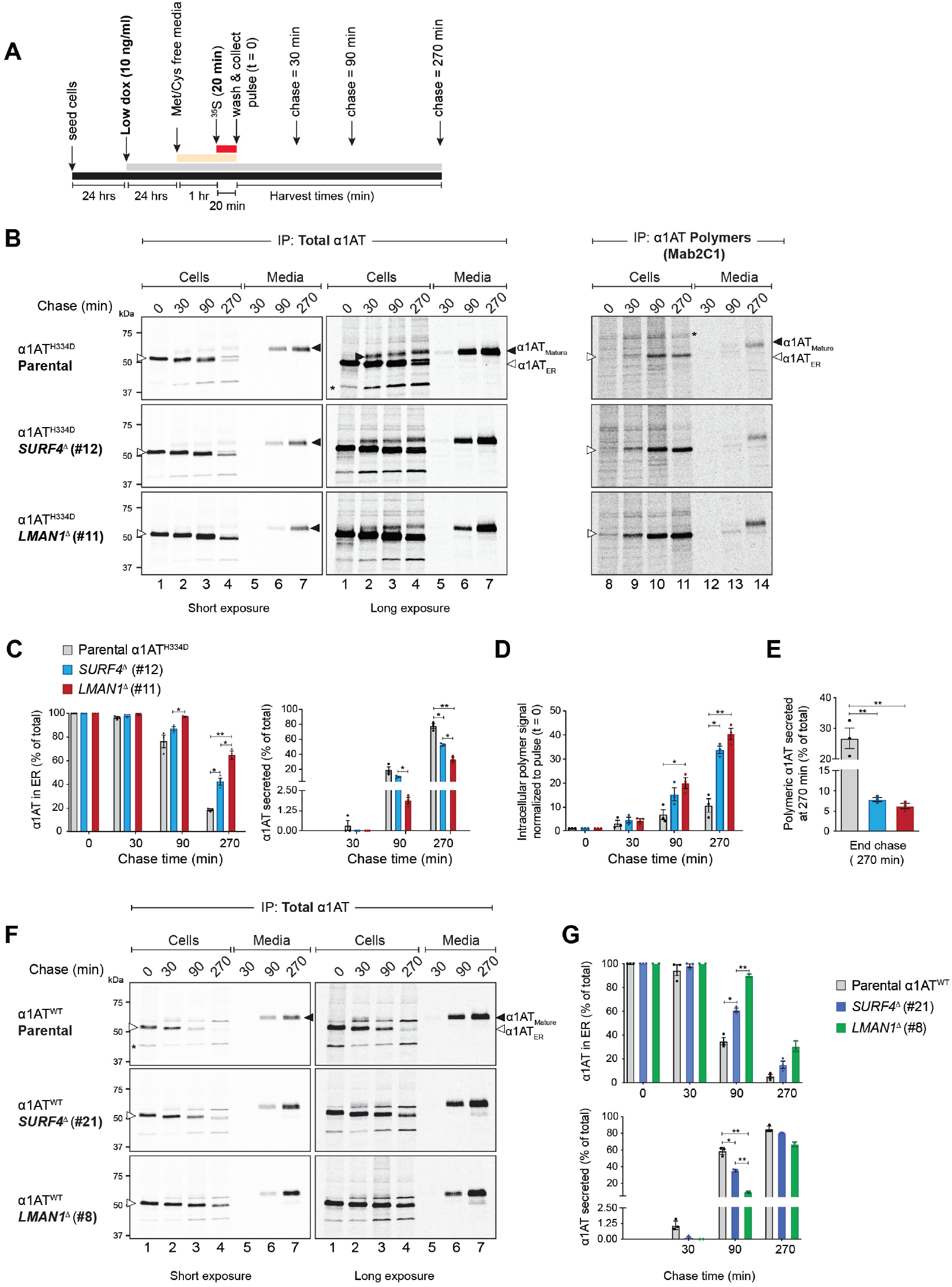
Altered intracellular trafficking of α1-antitrypsin in *SURF4* and *LMAN1* disrupted cells. **(A)** Schema of the experimental design. Note the induction of α1AT expression with low concentration (10 ng/ml) of doxycycline (dox), ^35^S-methionine/cysteine (Met/Cys) pulse labelling (20 min) and chase times (30-270 min). **(B)** Short and long exposures of autoradiographs of SDS-PAGE gels load with labelled α1AT immunoprecipitated with a polyclonal antibody reactive with all forms of α1AT (left panels) or Mab2C1, selective for α1AT polymers (right panel) from lysates of parental CHO-K1 Tet-on_α1AT^H334D^ cells and their *SURF4*^Δ^ and *LMAN1*^Δ^ derivatives (‘Cells’) or the culture supernatant (‘Media’). White arrowheads indicate the ER-associated form (α1AT_ER_) and black arrowheads the mature-glycosylated form (α1AT_Mature_). Asterisks (*) represent unspecific bands. **(C)** Percentage of α1AT^H334D^ retained in the ER [(α1AT_ER_ in ‘B’), left panel] or secreted into the media (right panel) of total protein [‘cell’ signal + ‘media’ signal] at each time point. **(D)** Intracellular polymer signal normalised to α1AT polymer signal at pulse end (lane 8). **(E)** Percentage of α1AT polymers present in the media of total protein at 270 min, calculated as in ‘C’. **(F)** As in ‘B’, but using parental CHO-K1 Tet-on_α1AT^WT^ cells and their *SURF4*^Δ^ and *LMAN1*^Δ^ derivatives. Total α1AT from cells and media was immunoprecipitated as in ‘B’. **(G)** Percentage of α1AT^WT^ retained in the ER (upper panel) or secreted into the media (lower panel), calculated as in ‘C’. Autoradiographs are representative of three independent experiments except for *LMAN1*^Δ^ (clone #8, n = 2). Quantitative plots show the mean ± SEM. *, P < 0.05; and **, P < 0.01. Two-way (in ‘C’, ‘D’ and ‘G’) or one-way ANOVA (in ‘E’) followed by Tukey’s post-hoc multiple comparison test.

Having confirmed a role for LMAN1 and SURF4 in trafficking of α1AT^H334D^, we then sought to determine their role in trafficking of α1AT^WT^ in CHO cells. The same pulse-chase labelling procedure described above was applied to parental, *SURF4*^Δ^ and *LMAN1*^Δ^ CHO-K1 Tet-on_α1AT^WT^ cells. Clearance of wild-type, monomeric α1AT from the ER was significantly delayed in *LMAN1*^Δ^ cells, consistent with previous observations (Nyfeler et al., 2008; Zhang et al., 2011), but also in *SURF4*^Δ^ cells, albeit to a lesser degree (Fig. 5F,G). Of note, the accumulation of wild-type monomer in *SURF4*^Δ^ and *LMAN1*^Δ^ cells did not result in detectable polymer formation by ELISA.

These observations implicate both LMAN1 and SURF4 in trafficking of wild-type and polymerogenic α1AT in CHO-K1 cells. This explains enhanced intracellular accumulation of α1AT polymers observed in the α1AT^H334D^-expressing cells lacking either LMAN1 or SURF4.

### SURF4 disruption preferentially impairs intracellular trafficking of α1AT polymers

SURF4 has been proposed as an ER cargo receptor that prioritises export of large, polymeric proteins (Saegusa et al., 2018; Yin et al., 2018). This, together with our observations noted above, suggested the possibility that SURF4 might also have a role in facilitating the exit of α1AT polymers from the ER. To address this question, we modified the pulse-chase procedure: synthesis of α1AT^H334D^ was increased by treating the cells with a higher concentration of doxycycline, thus shifting the equilibrium towards polymer formation. Crucially, the pulse and chase windows were prolonged to allow clearance of the fast-trafficking (labelled) mutant monomeric species and thereby focused the analysis on the remaining polymers (Fig. 6A).

**Fig. 6.**
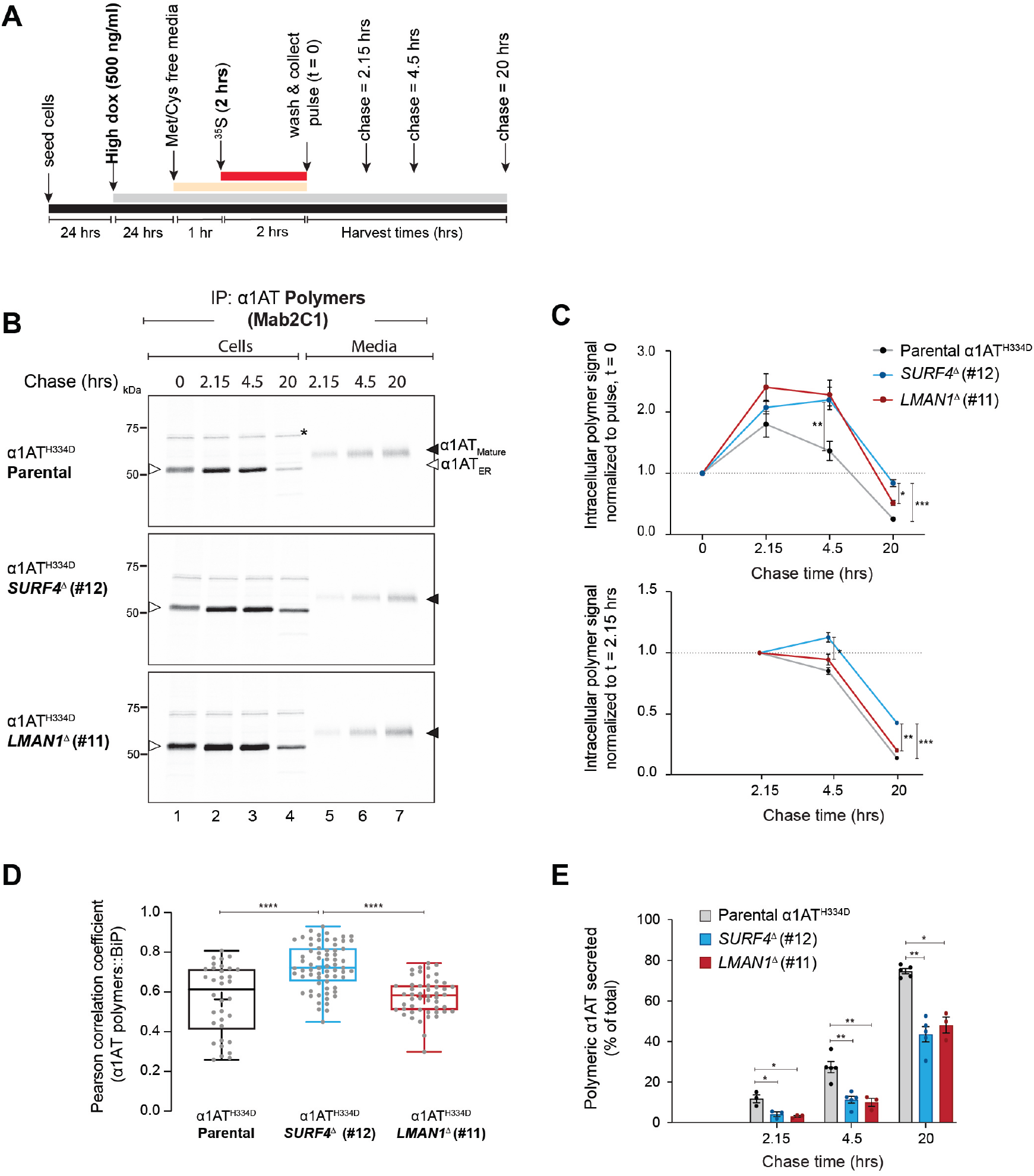
SURF4 and LMAN1 favour ER exit of α1-antitrypsin polymers. **(A)** Schema of the experimental design. Note the induction of α1AT expression with a high concentration (500 ng/ml) of doxycycline (dox) and the lengthy ^35^S-Met/Cys pulse labelling period (2 hrs) and chase times (2.15 - 20 hrs). **(B)** Autoradiographs of SDS-PAGE gels loaded with labelled α1AT immunoprecipitated with polymer-selective Mab2C1 from lysates of parental CHO-K1 Tet-on_α1AT^H334D^ cells and their *SURF4*^Δ^ and *LMAN1*^Δ^ derivatives (‘Cells’) or the culture supernatant (‘Media’). White arrowheads indicate the ER-associated form (α1AT_ER_) and black arrowheads the mature-glycosylated form (α1AT_Mature_). Asterisks (*) represent unspecific bands. **(C)** Plot of the cell-associated α1AT polymer signal at the indicated times, normalised to the signal at pulse end (lane 1; upper panel) or to the signal at 2.15 hrs (lane 2; bottom panel). **(D)** Pearson coefficient for the co-localisation of α1AT polymers (Mab2C1-stained) with the ER marker BiP in doxycycline-induced parental CHO-K1 Tet-on_α1AT^H334D^ cells (n = 34) and their *SURF4*^Δ^ (n = 67) and *LMAN1*^Δ^ (n = 50) derivatives (Fig. S4B). **(E)** Percentage of α1AT polymers present in the media of total protein [‘cell’ signal + ‘media’ signal] at each time point in ‘B’. Quantitative plots show the mean ± SEM (n = 3-5). *, P < 0.05; **, P < 0.01; ***, P < 0.001; and ***, P < 0.0001. Two-way (in ‘C’ and ‘E’) or one-way ANOVA (in ‘D’) followed by Tukey’s post-hoc multiple comparison test

The efficacy of these modifications is reflected in the appearance of a detectable pool of intracellular polymers at the end of the pulse and their persistence throughout the lengthy chase period, more conspicuously so in the *SURF4*^Δ^ and *LMAN1*^Δ^ cells (Fig. 6B). In all three genotypes, labelled polymers also appeared in the culture media (Fig. 6B) and these exhibited slower mobility on SDS-PAGE, compared to the cell-associated polymers. This observation is consistent with post-ER glycan modifications and indicates conventional trafficking through the secretory pathway.

In all three genotypes, intracellular polymer levels continued to increase after the pulse with levels peaking between 2.15-4.5 hrs chase (Fig. 6C, upper panel). Thus, considering this peak as a reference point by which to track the fate of ER-localised polymers, we found that *SURF4*^Δ^ cells retained proportionally more polymers compared to parental or *LMAN1*^Δ^ cells (Fig. 6C, lower panel). This finding correlated with higher degree of co-localisation of the polymers with the ER marker BiP in *SURF4*^Δ^ cells (Fig. 6D and Fig. S4B). Notably, the kinetics of the ratio of secreted polymers to cell-associated polymers was significantly slower in *LMAN1*^Δ^ and *SURF4*^Δ^ cells (Fig. 6E). Similar results were obtained with another independently derived *SURF4*^Δ^ clone (Fig. S4).

These findings implicate both LMAN1 and SURF4 in secretion of α1AT polymers in CHO-K1 cells and suggest a preference of SURF4 for the transport of intracellular α1AT polymers out of the ER compared to LMAN1.

### SURF4 interacts with α1AT in CHO-K1 cells

The interaction of LMAN1 and α1AT has been previously explored (Nyfeler et al., 2008). To assess possible physical interactions of SURF4 and α1AT, cells expressing α1AT^H334D^ or α1AT^WT^ were transfected with FLAG-tagged SURF4 and subjected to crosslinked. FLAG-tagged SURF4 was selectively recovered by anti-FLAG immunoprecipitation, accompanied by either α1AT^WT^ or α1AT^H334D^ (Fig. 7A,B). Transfection with a 7xHis-tagged SURF4 provided an opportunity to recover SURF4-α1AT complexes under denaturing conditions, which also allowed more stringent wash steps. Nickel affinity pulldowns indicated that both α1AT^WT^ and α1AT^H334D^ were recovered in complex with 7xHis-SURF4 (Fig. 7C,D). Their recovery under denaturing conditions is consistent with a proximal interaction between the two species, though bridging by a third factor cannot be excluded.

**Fig. 7.**
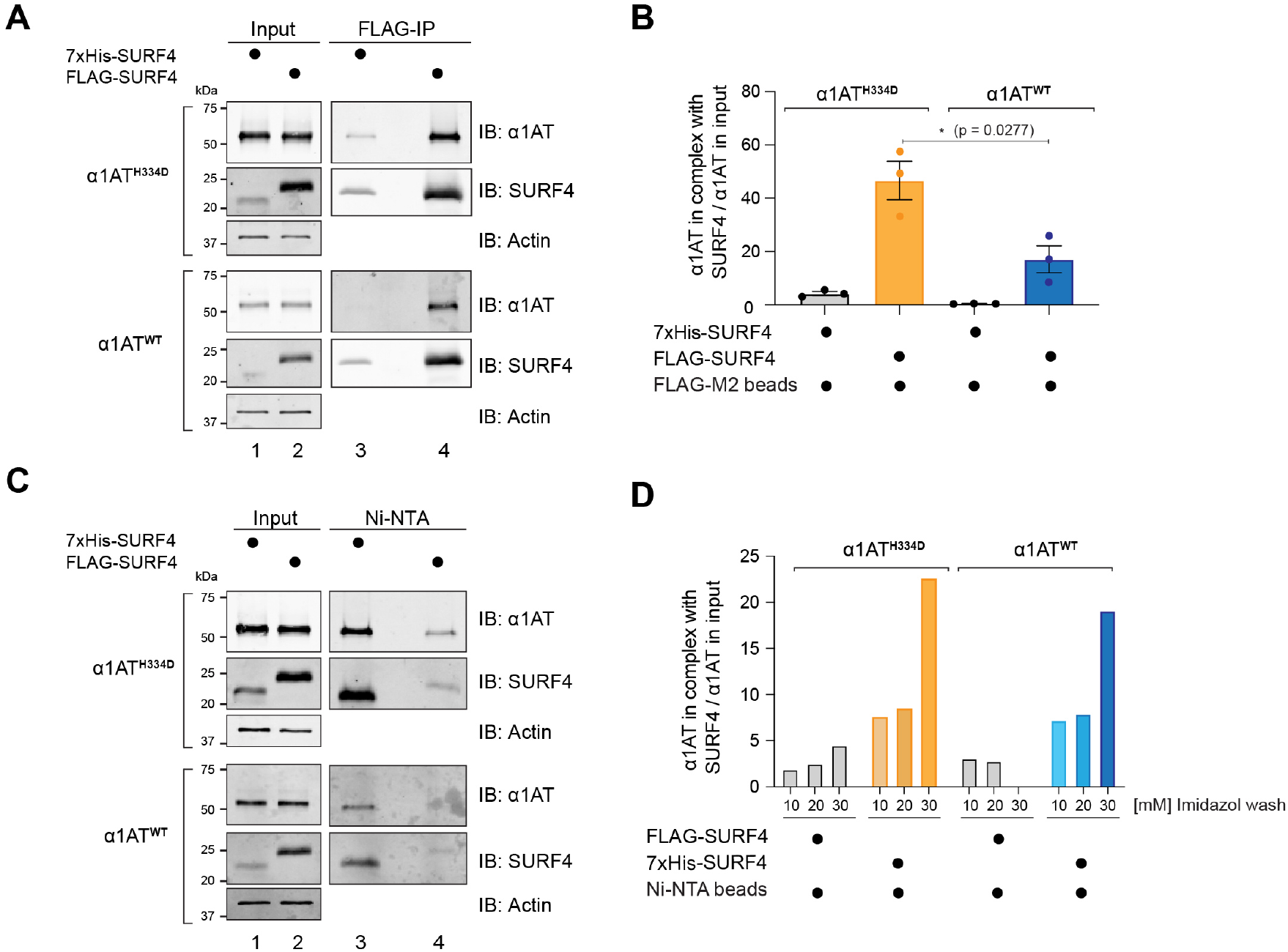
SURF4 interacts with α1-antitrypsin. **(A)** Representative immunoblots of α1AT recovered in complex with FLAG-SURF4 (FLAG-IP) from CHO-K1 Tet-on cells expressing α1AT^WT^ or α1AT^H334D^ transfected with a FLAG-tagged or 7xHis-tagged (as control) SURF4 plasmids and subjected to crosslinking. **(B)** Ratio of the signal from the α1AT recovered in complex with FLAG-SURF4 to the α1AT signal in the ‘input’. Shown is mean ± SEM from three independent experiments as in ‘A’ (Student’s t test). **(C)** As in ‘A’, but performing Ni-NTA affinity pulldowns under denaturing conditions on the same lysates used in ‘A’. An imidazole gradient from 10-30 mM in the wash buffer was used across three experiments. This SDS-PAGE gel represents samples washed with 30 mM imidazole. Cells transfected with a FLAG-tagged SURF4 reported on the background in this assay. **(D)** Ratio of the signal from the α1AT recovered in complex with 7xHis-tagged SURF4 to the α1AT signal in the ‘input’ from three different experiments performed as in ‘C’ in buffers with the indicated concentration of imidazole.

The evidence provided here for an interaction between SURF4 and α1AT is in keeping with SURF4’s functional role in trafficking of both polymeric and monomeric forms of α1AT.

## Discussion

By interfering with secretion, intracellular polymerisation of mutant α1AT limits its plasma concentration and contributes to the loss-of-function features of α1AT deficiency. Simultaneously, polymer retention contributes to gain-of-function features such as liver cirrhosis, whilst extracellular polymers appear to play a pro-inflammatory role in the lung (Lomas and Mahadeva, 2002) and elsewhere (Gross et al., 2009; Morris et al., 2011). Here, an unbiased genome-wide screen identified modifiers of intracellular levels of α1AT polymers, uncovering a previously under-appreciated role for cargo receptors in their active export from the ER and ultimately secretion of a fraction of the intracellular pool.

The strongest coherent signature to emerge from our screen was factors involved in cargo exit from the ER. These included LMAN1, a transmembrane cargo receptor known to have a role in the ER export of wild-type α1AT (Nyfeler et al., 2008; Zhang et al., 2011), validating the experimental approach. The screen also implicated SURF4 in affecting the intracellular levels of α1AT polymers. SURF4, the human orthologue of the yeast cargo receptor Erv29p (Belden and Barlowe, 2001), has been shown to be a versatile multi-spanning cargo receptor that facilitates export of large proteins such as the 550-kDa apolipoprotein B (Saegusa et al., 2018), small proteins such as the 75-kDa PCSK9 (Emmer et al., 2018), and soluble cargos that tend to aggregate within the ER (Yin et al., 2018). SURF4 has not been previously-recognised to have a role in the trafficking of α1AT, but it has been reported to form multiprotein complexes with LMAN1, along with other components of the ER exit complex (Mitrovic et al., 2008). Therefore, we focused our attention on the mechanisms by which loss of these cargo receptors altered the intracellular fate of α1AT. These studies were carried out in genetically malleable CHO-K1 cells that recapitulate both ER morphology changes observed in hepatocytes of α1AT-deficient patients (Ordonez et al., 2013) and the impairment of intracellular protein mobility observed in induced pluripotent stem cell-derived α1AT deficiency hepatocytes (Segeritz et al., 2018).

Disruption of either *LMAN1* or *SURF4* delayed trafficking of both polymerogenic α1AT^H334D^ and α1AT^WT^ out of the ER in this CHO-K1 system. As polymerisation is a concentration-dependent process (Lomas et al., 1993), impaired ER egress of mutant α1AT monomers could account for all the increase in intracellular polymer signal observed in the *LMAN1*^*Δ*^ and *SURF4*^*Δ*^ cells. This finding nonetheless emphasises the fact that variation in the efficiency of monomer trafficking out of the ER could contribute to the clinical heterogeneity in polymer-induced liver disease (Wu et al., 1994).

Less anticipated were findings pointing to a role for LMAN1 and SURF4 in the egress of polymers out of the ER and, ultimately, in their secretion from cells. This insight was gleaned from cells expressing high levels of mutant α1AT^H334D^, conditions predicted to shift the equilibrium in the ER towards polymerisation. Introducing a delay in the pulse-chase experiment that favoured clearance of residual fast-trafficking labelled monomers, focused the analysis on the fate of polymers. *LMAN1*^Δ^ and even more so *SURF4*^Δ^ cells retained relatively more polymers and secreted relatively fewer polymers than parental cells. Co-localisation of the excess polymers with the ER marker BiP, was particularly conspicuous in the *SURF4*^Δ^ cells, supporting the idea that SURF4 may have an important role in clearing the ER of α1AT polymers and possibly other large cargos, as suggested previously (Saegusa et al., 2018).

Co-immunoprecipitation experiments hinted at direct contact, or at least close proximity between SURF4 and α1AT. This was observed despite the absence from α1AT of an N-terminal motif previously reported to promote cargo binding to SURF4 (Yin et al., 2018) but also absent from other putative SURF4 cargos (e.g., PCSK9 and apolipoprotein B). Thus, at present, the basis for SURF4’s ability to select monomeric and polymeric α1AT for export from the ER remains unknown.

The mobility of α1AT during SDS-PAGE suggests that polymeric α1AT found in the culture supernatant had undergone post-ER glycan modifications. This finding, together with the genetic evidence of a role for ER cargo receptors in its itinerary suggests that at least a fraction of extracellular polymers found their way through the conventional secretory pathway. The existence of pathway(s) by which misfolded ER proteins traffic out of the compartment, ultimately to be degraded in the lysosome (Fregno et al., 2018), raises the possibility that LMAN1 or SURF4 also restrain intracellular polymer levels by promoting a trafficking event that contributes to their intracellular degradation. These issues remain unsettled even in our CHO-K1 model. Nonetheless, the role of ER cargo receptors in the itinerary of α1AT monomers and polymers highlighted in this study conjures the possibility of mechanism-based interventions to alter the balance of polymers retained in cells, degraded intracellularly or secreted and could represent new therapeutic targets for the underlying lung disease.

## Acknowledgments

We thank the CIMR flow cytometry (Reiner Schulte, Chiara Cossetti and Gabriela Grondys-Kotarba) and microscopy teams (Matthew Gratian and Mark Bowen) for technical support, Marcella Ma and Brian Lam (CRUK) for assistance with NGS and our lab members, especially Steffen Preissler (CIMR) for critical comments and advice.

## Author’s contributions

**A**.**O**. conceived, initiated, and led the project, designed and conducted the experiments, analysed and interpreted the data, prepared figures and tables and wrote the first draft of the manuscript. **H**.**P**.**H**. designed the CHO CRISPR/Cas9 library and contributed experimentally with the lentiviral library, in data analysis and reviewed the manuscript. **S**.**J**.**M**. contributed to discussion and revision of the manuscript. **D**.**R**. conceived and oversaw the project, interpreted the data, and co-wrote the manuscript. All authors read and approved the final manuscript.

## Material and methods

Standard molecular cloning methods were used to create the plasmids DNAs listed in the Table S2. Single guides RNAs (sgRNAs) and oligonucleotides are listed in Table S3. Antibodies, reagents and software are listed in Table S4.

### Cell culture

CHO-K1 cells expressing human α1AT^WT^ or the polymerogenic α1AT^H334D^ mutant under a tetracycline inducible promoter (Ordonez et al., 2013) were cultured in DMEM cell media (Sigma) supplemented with 10% Tet-free serum (Pan-Biotech), 1x Penicillin-Streptomycin (Sigma), 1x MEM-non-essential-amino-acids (Sigma), 2 mM L-glutamine (Sigma), 200 μg/mL G418 and 500 μg/mL of Hygromycin B (Invitrogen). Depending on the experiment, α1AT expression was induced with 10 ng/ml (‘low dox’) or 500 ng/ml (‘high dox’) doxycycline for 24 hrs. Although not relevant for these experiments, the open reading frame of *Cricetulus griseus DDIT3* locus was replaced by GFP (*CHOP::GFP* reporter) in the parental CHO-K1 Tet-on cells. For the CRISPR/Cas9 screen we stably introduced the Cas9 nuclease into CHO-K1 Tet-on_α1AT^H334D^ cells via lentiviral transduction (UK1714, see Table S2 and Table S3). Cas9 activity in derivative cell lines was confirmed by targeting the *CHOP::GFP* reporter with a EGFP-targeting sgRNA (UK1717) followed by induction of ER stress.

CHO-K1 S21 cells bearing *CHOP::GFP* and *XBP1s::Turquoise* reporters (Sekine et al., 2016) were maintained in Nutrient Mixture F12 (Sigma) supplemented with 10% serum (FetalClone II, ThermoScientific), 1× Penicillin-Streptomycin and 2 mM L-glutamine. HEK293T cells (ATCC CRL-3216) were cultured in DMEM supplemented as above. All cells were grown at 37°C and 5% CO_2_.

### Lentivirus production

Lentiviral particles were produced by transfecting HEK293T cells with the library plasmids (UK2561, UK2321 and UK2378) together with the packaging plasmids psPAX2 (UK1701) and pMD2.G (UK1700) at a 10:7.5:5 ratio using TransIT-293 reagent (Mirus). The supernatant containing the viral particles was collected 48 hrs after transfection, filtered through a 0.45 μm filter, and directly used to infect CHO-K1 cells seeded in 6-well plates for viral titration.

### Intracellular polymer staining, FACS and flow cytometry

Cells were washed twice with PBS, collected in PBS containing 4 mM EDTA and 0.2% BSA and fixed in 1% formaldehyde for 10 min. Fixative was washed-out at 700× *g* for 5 min and cells were permeabilised in blocking buffer [PBS containing 0.1% Triton X-100 and 10% FBS] for 20 min, incubated with the primary α1AT polymer-specific monoclonal antibody 2C1 (Mab2C1) (Miranda et al., 2010) for 30 min, washed three times in blocking solution, and then incubated with the secondary DyLight 633-labelled anti-mouse antibody for 20 min. Cells were washed, resuspended in PBS containing 2 mM EDTA and 2% FBS, filtered and sorted on an Influx cell sorter (BD) or analysed by flow cytometry (20,000 cells/sample) using a LSRFortessa cell analyser (BD). In order to reduce cell clumping, a cell density of ∼2×10^6^ cells/ml was adjusted and all incubations were done with orbital agitation at room temperature or 4°C, when required. α1AT polymers (Mab2C1 signal) were detected by excitation at 640 nm and monitoring emission at 670/14 nm; blue fluorescent protein (BFP) by excitation at 405 nm and monitoring at 450/50 nm; *CHOP::GFP* by excitation at 488 nm and monitoring at 530/30 nm; *XBP1s::Turquoise* by excitation at 405 nm and monitoring at 450/50 nm. Data were processed using FlowJo and statistical analysis using Prism8 (GraphPad).

The sensitivity to UPR induction in CHO-K1 S21 cells bearing *CHOP::GFP* and *XBP1s::Turquoise* reporters was analysed after transient transfection with 1 μg sgRNA-mCherry-Cas9 encoding plasmids, targeting *SURF4, LMAN1* and *HSPA5* (BiP protein) (see Table S2). Each gene was targeted with two different sgRNA and four days after transfection cells were analysed by flow cytometry.

### Whole genome CRISPR screen

High-throughput screen was carried out as previously described (Shalem et al., 2014) using a Chinese hamster knockout CRISPR/Cas9 library containing 125,030 sgRNAs targeting 20,680 genes (most with 6 guides per gene) as well as 1,239 non-targeting sgRNAs as a negative control cloned into the lentiviral sgRNA expression vector pKLV-U6gRNA(BbsI)-PGKpuro2ABFP as described (Harding et al., manuscript in preparation). Approximately 2.1×10^8^ CHO-K1 Tet-on_α1AT^H334D^_Cas9 cells were infected at a multiplicity of infection (MOI) of 0.3, to favour infection with a single viral particle/cell and selected with 8 μg/ml puromycin for 7 days. Expression of α1AT was induced with 10 ng/ml doxycycline for 24 hrs. Afterwards, the cells were fixed and permeabilised for intracellular staining of α1AT polymers. Approximately 6.6×10^7^ Mab2C1-stained fixed cells were subjected to FACS and collected in 3 bins according to their fluorescence intensity at 670 nm (Mab2C1): ‘brightest’ (∼2% of total sorted), ‘medium-bright’ (∼4.5% of total), and ‘dull’ (∼10% of total) as shown in Fig. 1B. Rounds of enrichment were carried on by extracting the genomic DNA of the ‘brightest’-binned fixed cells and recovering by PCR a 220bp fragment containing the sgRNA-bearing region (oligonucleotides 2182 and 1758). The amplicon was ligated into the parental lentiviral backbone (UK1789) to generate derivative enriched libraries (called Lib_1_ and Lib_2_) that were used to perform two successive cycles of infection of ∼2×10^7^ parental CHO-K1 Tet-on_α1AT^H334D^_Cas9 cells. In each round an equal number of infected, untreated cells (no doxycycline) or uninfected, doxycycline-treated cells were passed without sorting as a control group.

Genomic DNA from fixed, enriched, and sorted populations as well as fixed, unsorted libraries was extracted from ∼1-3×10^6^ and ∼3.6×10^7^ cells respectively, by incubation in proteinase K solution [100 mM Tris-HCl pH 8.5, 5 mM EDTA, 200 mM NaCl, 0.25% SDS, 0.2 mg/ml Proteinase K] overnight at 50°C. To reverse formaldehyde crosslinks, samples were supplemented with 500 mM NaCl and incubated at 65°C for 16 hrs. Integrated sgRNA sequences were amplified by nested PCR and the adaptors for Illumina sequencing (HiSeq4000) were introduced at the final amplification round using oligonucleotides 1759-1769 (Table S3). Downstream analysis to obtain sgRNA read counts, gene rankings, and statistics were obtained using the MAGeCK computational software (Li et al., 2014). Gene ontology analyses were performed using Metascape software with default parameters (Zhou et al., 2019).

### Validation of candidate genes

Two individual sgRNAs designed in the library targeting exon regions of *Cricetulus griseus LMAN1, SURF4* and *SEC23B* were cloned into the pSpCas9(BB)-2A-mCherry plasmid (UK1610) as previously reported (Ran et al., 2013). Cells were transfected with 1 μg of sgRNA/Cas9 plasmids UK2501-UK2506) using Lipofectamine LTX (Thermofisher). Forty-eight hours after transfection, mCherry-positive cells were individually sorted into 96-well plates using a MoFlo Cell Sorter (Beckman Coulter). The presence of frameshift-causing insertion/deletions in both alleles of the obtained clones was achieved by capillary electrophoresis on a 3730xl DNA analyser (Applied Biosystems) and amplifying the targeted region by PCR using a gene-specific 5′ 6-carboxyfluorescein (FAM)-labelled oligonucleotides (Hjelm et al., 2010). The knockouts were confirmed by Sanger sequencing and immunoblotting. Genomic information of the clones used in this study is provided in Table S5.

### Mammalian cell lysates, sandwich ELISA, and immunoblotting

Cells were lysed in Nonidet lysis buffer [150 mM NaCl, 50 mM Tris-HCl pH 7.5, 1% Nonidet P-40] supplemented with protease inhibitor mixture (Roche) for 20 min on ice. To quantify polymer and total levels of intracellular α1AT, cell lysates were analysed by sandwich ELISA using the polymer-specific Mab2C1 and a monoclonal antibody that recognises all α1AT conformers (Mab3C11) (Tan et al., 2015) respectively. Briefly, high binding surface COSTAR 96-well plates (Corning) were coated overnight with purified rabbit polyclonal antibody against total α1AT at 2 μg/ml in PBS. After washing with PBS containing 0.9% NaCl and 0.05% Tween-20, the plates were blocked for 1 hr in blocking buffer (PBS containing 0.25% BSA and 0.05% Tween-20). Samples and standard curves were diluted in blocking buffer and incubated for 2 hrs with the primary antibodies, Mab2C1 or Mab3C11. Anti-mouse IgG horseradish peroxidase-labelled antibody was used as a secondary antibody and incubated for 1 hr. The reaction was developed with TMB liquid substrate for 10 min in the dark, and the reaction was stopped with 1 M H_2_SO_4_. Absorbance was read at 450 nm on a microplate reader. For immunoblots, SDS sample buffer was added to the lysates and proteins were denatured by heating at 70°C for 10 min and separated on 10-12% SDS-PAGE gels and transferred onto PVDF membranes prior to immunodetection. Cyclophilin B and actin were detected as loading controls. To detect the multi-pass transmembrane protein SURF4, samples were incubated at 37°C for 15 min. Native-PAGE (4.5% stacking gel and a 7.5% separation gel) was performed to separate and identify α1AT monomers and polymers. Membranes were scanned using an Odyssey near infrared imager (LI-COR) and signals were quantified with ImageJ software.

### [^35^S] metabolic labelling and immunoprecipitation

Cells were starved in Methionine/Cysteine-free DMEM for 1 hr, pulsed with 100 μCi/well [^35^S]methionine/cysteine (Expre^35^S Protein Labelling Mix) and harvested or chased in DMEM containing 200 mM methionine and cysteine and 10% dialysed FBS. After the chase, culture media were collected and cells harvested on ice in Nonidet lysis buffer supplemented with protease inhibitor mixture (Roche). Culture media and cell lysates were precleared and α1AT was immunoprecipitated with a α1AT polyclonal antibody (total) or the Mab2C1 (polymer-specific) by splitting each sample in two equal parts. Radiolabelled proteins were recovered in 2×SDS-PAGE loading buffer, separated on 10% SDS-PAGE gels, detected by autoradiography with a Typhoon biomolecular imager (GE Healthcare) and quantified using ImageJ.

### Cross-linking and co-immunoprecipitation

CHO-K1 Tet-on cells expressing α1AT^WT^ or α1AT^H334D^ were grown in 10-cm dishes and transfected with either a 7×His- or FLAG-tagged SURF4 (UK2622 and UK2549) for 6 hrs. Afterwards, medium was exchanged against medium supplemented with 500 ng/ml doxycycline and cells were further incubated for 20 hrs. Cross-linking was performed following a previously-published protocol (Zlatic et al., 2010) with modifications. Cells were washed twice with PBS/Ca/Mg solution (PBS containing 0.1 mM CaCl_2_ and 1 mM MgCl_2_) and incubated for 2 hrs on ice with 1 mM dithiobis(succinimidyl propionate) (DSP, reversible crosslinker) diluted in pre-warmed (37°C) PBS/Ca/Mg solution. The DSP-containing solution was removed and the residual DSP was quenched for 15 min with PBS/Ca/Mg solution supplemented with 20 mM Tris-HCl pH 7.4. Cells were washed with PBS/Ca/Mg and lysed in Nonidet lysis buffer. A post-nuclear supernatant was prepared by centrifugation at 20,000× *g* at 4°C for 15 min, and then cleared again at 20,000× *g* for 5 min. For immunoprecipitation of FLAG-SURF4, cell lysates (750 μg total protein) were precleared with empty agarose beads and then incubated with anti-FLAG-M2 agarose affinity beads (Sigma) with rotation overnight at 4°C. Beads were washed four times with RIPA buffer [50 mM Tris-HCl pH 8, 150 mM NaCl, 1% triton X-100, 0.5% sodium deoxycholate, 0.1% SDS]. Bound proteins were eluted by addition of 2×SDS sample buffer (without DTT) and shacking at 37°C for 15 min to avoid aggregation of SURF4. Eluted proteins were recovered at 2,800× *g* for 5 min, 50 mM DTT was added and samples were further incubated at 37°C for 10 min. For pulldowns of 7xHis-SURF4, cell lysates were incubated in denaturing binding buffer (8 M Urea, 10 mM imidazole) containing protease inhibitors. Cell lysates were loaded onto Ni-NTA agarose beads (Qiagen) and incubated with orbital rotation overnight at RT. The beads were washed in denaturing washing buffer containing 150 mM NaCl, 50 mM Tris, 8 M Urea. Over the three independent experiments different concentrations of imidaole were used (10, 20 and 30 mM, respectively) to successively increase stringency of the wash step. Beads were then suspended in elution buffer [8 M Urea, 2% SDS, 50 mM DTT, 4 mM EDTA]. Equal volumes of the samples were loaded on 12% SDS-PAGE gels. Samples of the normalised cell lysates (15 μg) were loaded as ‘input’ controls and bands were quantitated using ImageJ.

### Confocal microscopy

Cells were seeded on coverslips pretreated with 0.1 mg/ml poly-L-lysine (Sigma) in 12-well plates and then fixed with 4% paraformaldehyde for 30 min, followed by permeabilisation with 0.1% Triton X-100 for 15 min. After 30 min blocking with PBS containing 10% BSA and 0.1% Triton X-100 the cells were co-stained with primary antibodies (Mab2C1 and anti-BiP) and the corresponding fluorescent secondary antibodies. Coverslips were mounted in FluorSave reagent (Calbiochem) containing 2% 1,4-diazabicyclo-[2.2.2]octane (Sigma). Imaging was performed on a Zeiss 710 confocal microscope using a 63x/1.4 oil immersion objective and diode, argon and HeNe lasers. The quantification of co-localisation between both fluorescence channels (Pearson correlation coefficient) was quantified using Volocity software, version 6.3 (PerkinElmer).

### Statistics

Data groups were analysed as described in the figure legends using GraphPad Prism8 software. Differences between groups were considered statistically significant if P < 0.05 (*, P < 0.05; **, P < 0.01; and ***, P < 0.001). All error bars represent mean ± SEM.

## Listing of supplemental materials

Fig. S1 shows the concentration-dependence of the response of CHO-K1 Tet-on cells to doxycycline and bafilomycinA1. Fig. S2 shows the quality control data analysis of the CRISPR/Cas9 screen performed by MAGeCK. Fig. S3 (related to Fig. 5) shows the altered intracellular trafficking of α1AT in an additional *SURF4*^Δ^ clone. Fig. S4 (related to Fig. 6) shows that SURF4 favours ER exit of α1AT polymers in an additional *SURF4*^Δ^ clone and contains microscopic images showing co-localisation of α1AT polymers with the ER marker BiP. Table S1 contains the complete ranked list of genes enriched in ‘brightest’ cells generated by MAGeCK. Table S2 indicates the recombinant plasmid DNAs used in this study. Table S3 indicates the list of sgRNAs and oligonucleotides. Table S4 indicates the list of antibodies, reagents and software. Table S5 indicated the clones generated in this study.

## Supplementary Figures

**Fig. S1.**
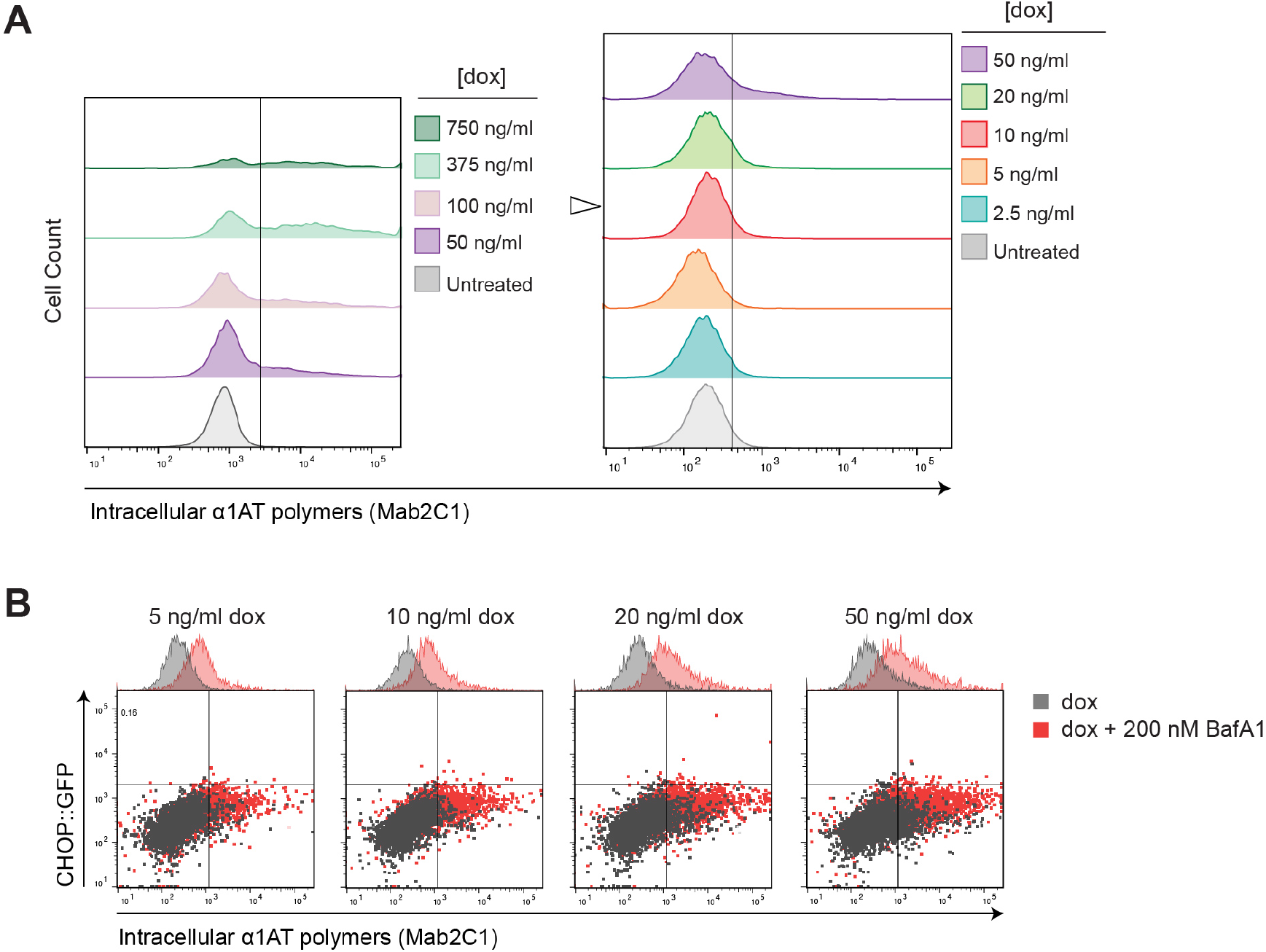
Concentration-dependence of the response of CHO-K1 Tet-on cells to doxycycline and bafilomycinA1. **(A)** Flow cytometry analysis of the fluorescence intensity as a measure of intracellular α1AT polymer levels (stained with Mab2C1) in CHO-K1 Tet-on_α1AT^H334D^_Cas9 cells treated for 24 hrs with the indicated concentrations of doxycycline (dox). The left and right panels represent two independent experiments. The white arrowhead indicates the dox concentration used in the screen. **(B)** Dual-channel flow cytometry of the UPR marker, *CHOP::GFP*, and intracellular levels of α1AT polymers in CHO-K1 Tet-on_α1AT^H334D^_Cas9 cells treated for 24 hrs with the indicated concentration of dox in presence or absence of bafilomycinA1 (BafA1; 200 nM, added during the last 16 hrs). 5,000 cells were analysed.

**Fig. S2.**
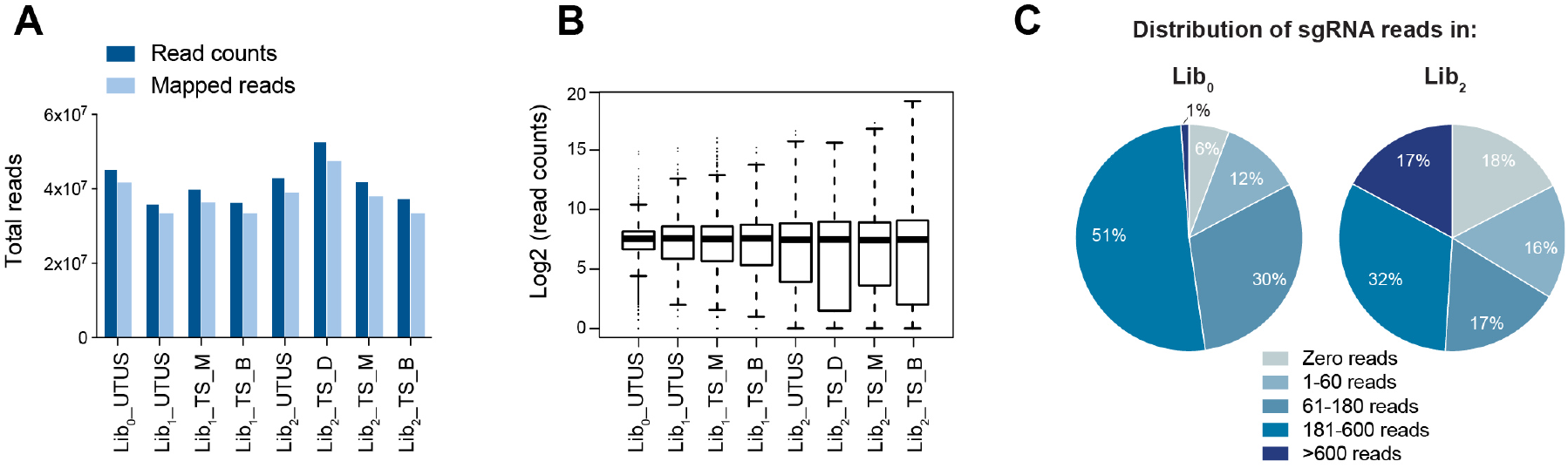
Quality control data analysis of the CRISPR/Cas9 screen performed by MAGeCK. **(A)** Total read counts and reads mapped to the CHO library analysed by MAGeCK [UTUS: untreated (no doxycycline) and unsorted; TS: treated (plus doxycycline) and sorted; Lib_0_: unenriched library) Lib_1_: derivative enriched library 1; Lib_2_: derivative enriched library 2; B: brightest; M: medium-bright; D: dull]. **(B)** Frequency distribution of sgRNA in each sample, showing the median-normalised read counts. **(C)** Representation of sgRNAs in unsorted cells after infection with the unenriched genome-wide library (Lib_0_) and enriched library (Lib_2_) according to their read counts.

**Fig. S3.**
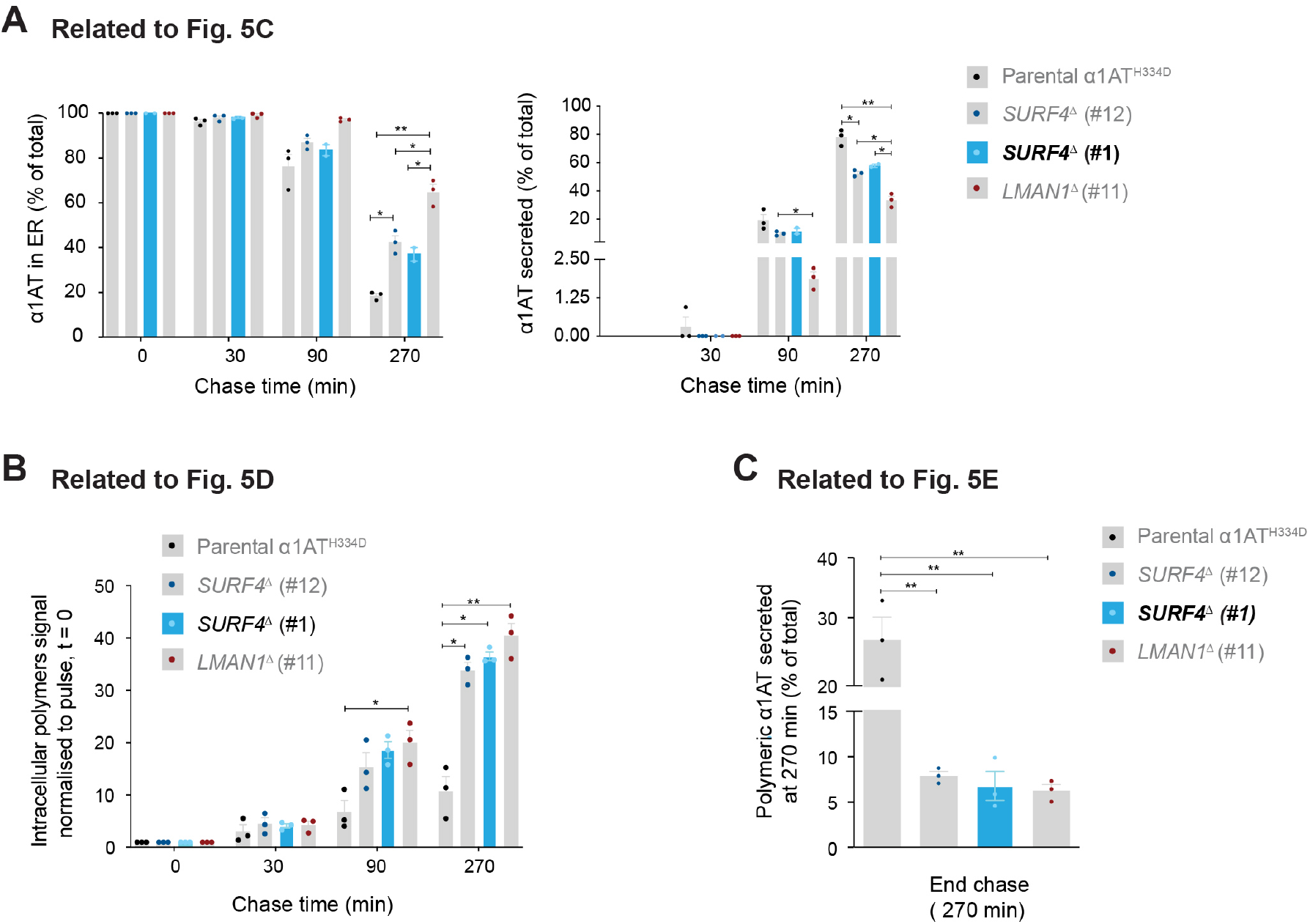
Altered intracellular trafficking of α1-antitrypsin in an additional *SURF4*^Δ^ clone. Labelled α1AT was immunoprecipitated with a polyclonal antibody reactive with all α1AT forms or a monoclonal antibody selective for α1AT polymers from lysates of parental CHO-K1 Tet-on_α1AT^H334D^ cells and their *SURF4*^Δ^ and *LMAN1*^Δ^ derivatives or from the culture media supernatant. **(A) Related to Fig. 5C**. Plots of the percentage of α1AT retained in the ER (left panel) or secreted into the media (right panel) at the indicated times. The additional *SURF4* disrupted clone [*SURF4*^Δ^ (#1)] is highlighted in blue and the other three genotypes (previously shown in Fig. 5C) are coloured in grey. **(B) Related to Fig. 5D**. Plot of the intracellular polymer signal normalised to polymer α1AT signal at pulse end (t = 0) at the indicated times. The additional *SURF4*^Δ^ (#1) clone is highlighted in blue. **(C) Related to Fig. 5E**. Plot of the percentage of α1AT polymers present in the media at 270 min. The additional *SURF4*^Δ^ (#1) clone is highlighted in blue. All quantitative plots show the mean ± SEM of two or three independent experiments; *p<0.05, **p<0.01. Two-way (in ‘A’ and ‘B’) or one-way ANOVA (in ‘C’) followed by Tukey’s post-hoc multiple comparison test.

**Fig. S4.**
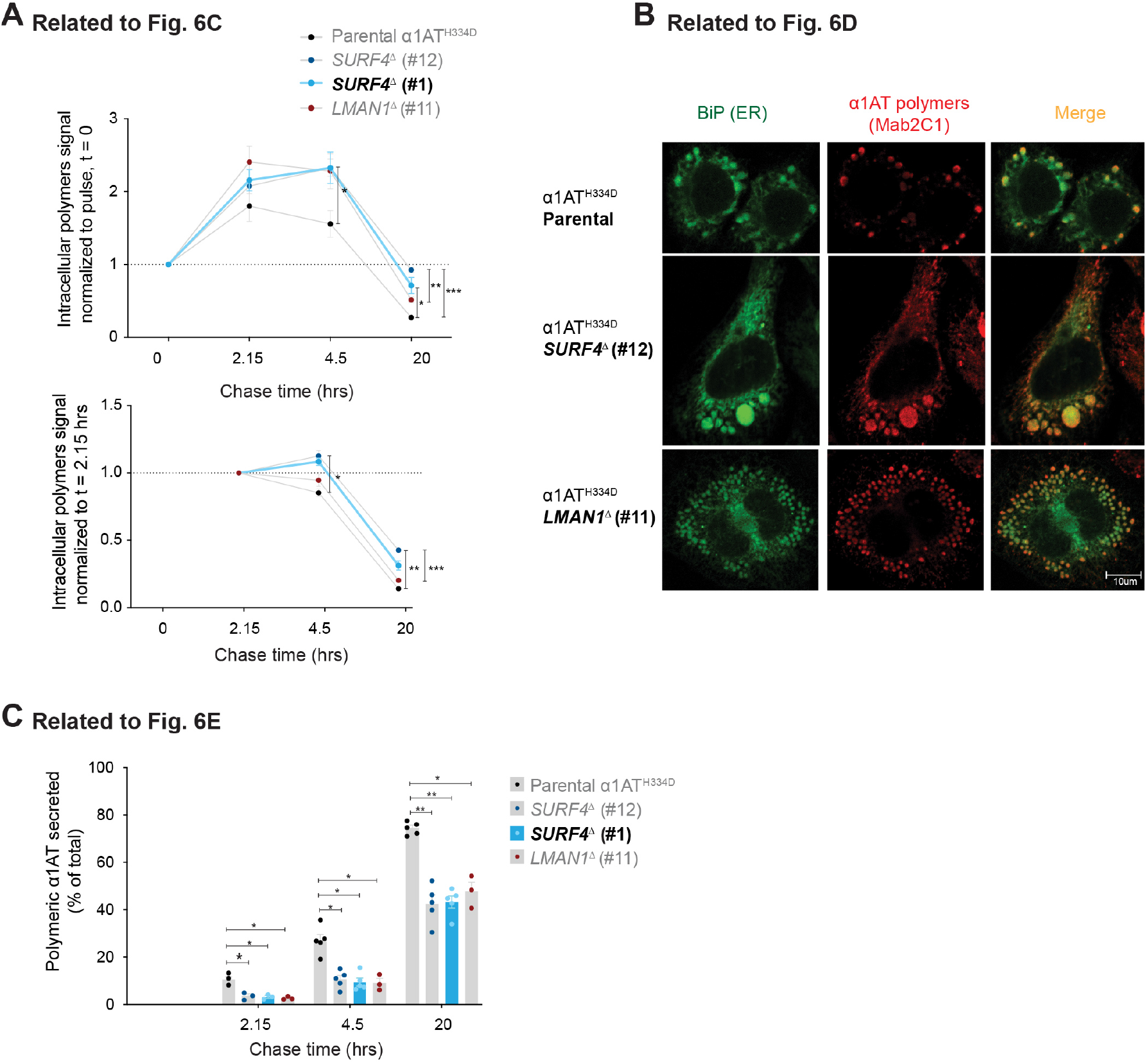
SURF4 favours ER exit of α1-antitrypsin polymers in an additional *SURF4*^Δ^ clone. Labelled α1AT was immunoprecipitated with a monoclonal antibody selective for α1AT polymers from lysates of parental CHO-K1 Tet-on_α1AT^H334D^ cells and their *SURF4*^Δ^ and *LMAN1*^Δ^ derivatives or from the culture media supernatant. **(A) Related to Fig. 6C**. Plots of the cell-associated α1AT polymer signal at the indicated times, normalised to the signal at pulse end [t = 0, (upper panel)] or at 2.15 hrs (bottom panel). The additional SURF4 disrupted clone [*SURF4*^Δ^ (#1)] is highlighted in blue and the other three genotypes (previously showed in Fig. 6C) are coloured in grey. **(B) Related to Fig. 6D**. Representative confocal immunofluorescence microscopy images of α1AT polymers (Mab2C1, red) together with an ER marker (BiP, green) in fixed parental CHO-K1 Tet-on_α1AT^H334D^ cells and their *SURF4*^Δ^ (#12) and *LMAN1*^Δ^ (#11) derivatives clones. α1AT expression was induced with 500 ng/ml doxycycline for 24 hrs. **(C) Related to Fig. 6E**. Percentage of α1AT polymers present in the media at the indicated times. The additional *SURF4*^Δ^ (#1) clone is highlighted in blue. All quantitative plots show the mean ± SEM of three to five independent experiments; *p<0.05, **p<0.01, ***p<0.001, ****p<0.0001. Two-way ANOVA test followed by Tukey’s post-hoc multiple comparison test.

## Supplementary Tables

**Table S1: Rank of genes enriched in cells with the highest level of polymer signal** (see attached file). Rank of genes enriched in cells infected with the “derivative enriched Lib_2_” doxycycline-treated and sorted, relative to cells infected with the “unenriched Lib_0_” untreated and unsorted. Genes are ranked by “pos | rank” with *ATF7IP* (activating transcription factor 7 interacting protein) on the top of positively selected genes. The top 121 positively selected genes correspond to Fig. 2A.

**Table S2:** Recombinant DNA used in this study.

| Lab ID | Plasmid name | Description | Reference |
| --- | --- | --- | --- |
| UK1610 | pSpCas9(BB)-2A-mCherry | Modified pSpCas9(BB)-2A vector to express mCherry together with guide RNA & Cas9 | Amin-Wetzel N et al., 2017 |
| UK1700 | pMD2.G | Addgene plasmid 12259, lentiviral packaging helper, (VSVG) | Unpublished, gift from Didier Trono |
| UK1701 | psPAX2 | Addgene plasmid 12260, next gen lentiviral packaging helper | Unpublished, gift from Didier Trono |
| UK1702 | LentiGuide-puro | Addgene plasmid 52963 | Sanjana et al., 2014, gift from Feng Zhang |
| UK1714 | Lenti-Cas9 | Lenti-Cas9 in which 2TA-blast sequence is removed to make a lenti-Cas9 without resistance selection marker | This study |
| UK1717 | EGFPsgRNA_lentiGuide-Puro | Lentiviral vector expressing EGFP CRISPR guides without expression of Cas9 | This study |
| UK1789 | pKLV-U6gRNA(BbsI)-PGKpuro2ABFP | Addgene 50946, BFP-2A-Puro tagged gRNAvector | Koike-Yusa et al., 2014, gift from Kosuke Yusa |
| UK1857 | cgHSPA5_g1_pSpCas(BB)-2A-mCherry | mCherry-tagged CRISPR plasmid (UK1610) for targeting hamster HSPA5 (BiP) | Preissler et al., 2017 |
| UK1858 | cgHSPA5_g2_pSpCas(BB)-2A-mCherry | mCherry-tagged CRISPR plasmid (UK1610) for targeting hamster HSPA5 (BiP) | Preissler et al., 2017 |
| UK2561 | pKLV-CHO_libA-PGKpuro2ABFP (Library0) | CHO CRISPR KO library of 125030 selected guides for whole genome CRISPR screening | Unpublished |
| UK2321 | pKLV-α1AT derivative enriched CHO library1 (MluI_BamHI)-PGKpuro2ABFP | CHO CRISPR KO derivative library 1 (Lib1) for α1AT polymer enrichment_Brightest population-After first sorting | This study |
| UK2378 | pKLV-α1AT derivative enriched CHO library2 (MluI_BamHI)-PGKpuro2ABFP | CHO CRISPR KO derivative library 2 (Lib2) for α1AT polymer enrichment_Brightest population-After second sorting | This study |
| UK2501 | cgLman1_g1_exon 11_pSpCas9(BB)-2A-mCherry | mCherry-tagged CRISPR plasmid (UK1610) targeting cgLMAN1_guide 1 | This study |
| UK2502 | cgLman1_g2_exon 9_pSpCas9(BB)-2A-mCherry | mCherry-tagged CRISPR plasmid (UK1610) targeting cgLMAN1_guide 2 | This study |
| UK2503 | cgSurf4_g1_exon 5_pSpCas9(BB)-2A-mCherry | mCherry-tagged CRISPR plasmid (UK1610) targeting cgSURF4_guide 1 | This study |
| UK2504 | cgSurf4_g2_exon 2_pSpCas9(BB)-2A-mCherry | mCherry-tagged CRISPR plasmid (UK1610) targeting cgSURF4_guide 2 | This study |
| UK2505 | cgSec23b_g1_exon 7_pSpCas9(BB)-2A-mCherry | mCherry-tagged CRISPR plasmid (UK1610) targeting cgSEC23b_guide 1 | This study |
| UK2506 | cgSec23b_g2_exon 13_pSpCas9(BB)-2A-mCherry | mCherry-tagged CRISPR plasmid (UK1610) targeting cgSEC23b_guide 2 | This study |
| UK2549 | FLAG-tagged SURF4 [pNLF-FLAG-SURF4-puro) | FLAG-tagged SURF4 [pNLF-FLAG-SURF4-puro) | Emmer et al., 2018, gift from David Ginsburg |
| UK2622 | pNLF-H7-SURF4-puro | Mammalian expression plasmid 7xHis N-term tagged SURF4 | This study |

**Table S3:**
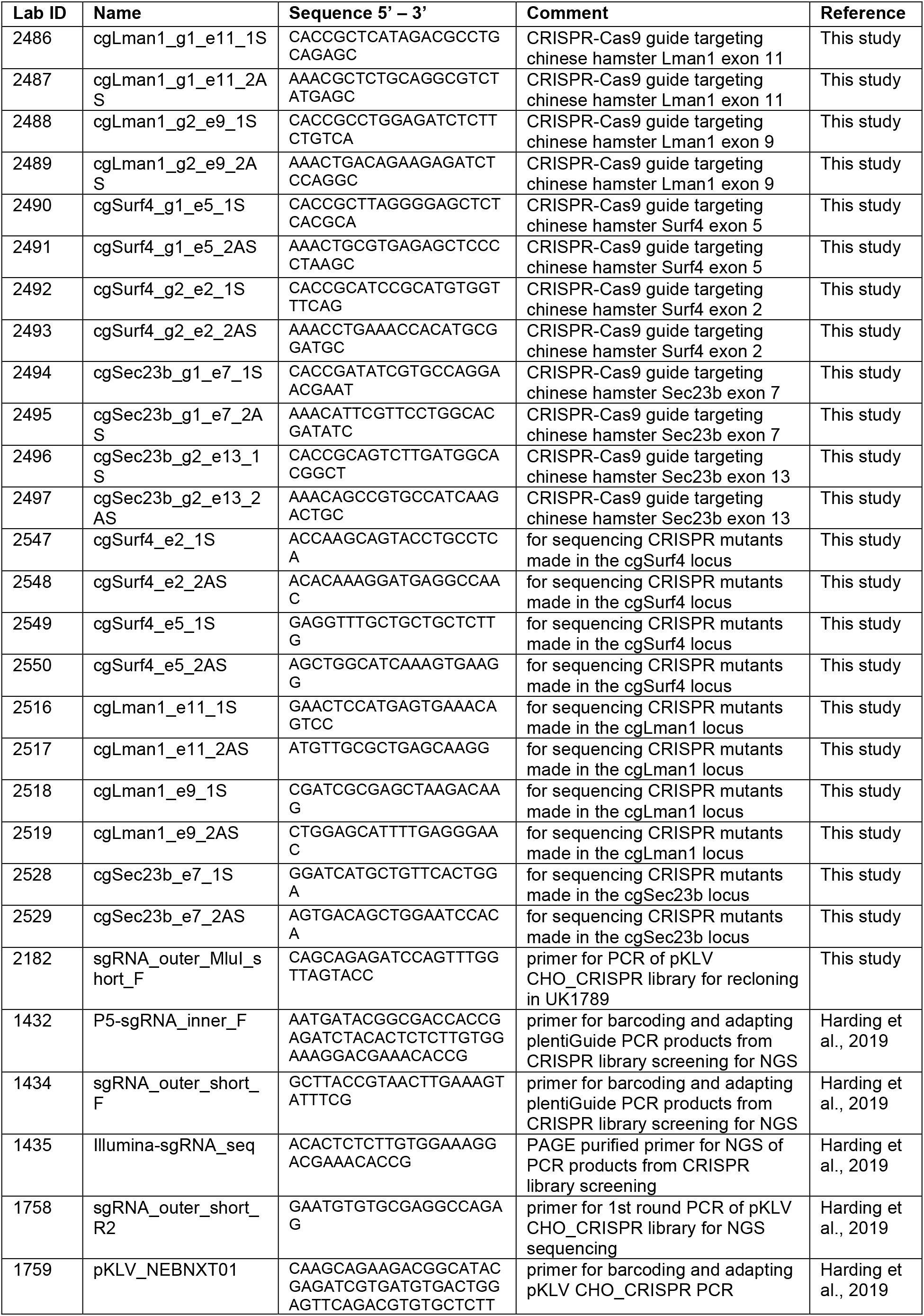
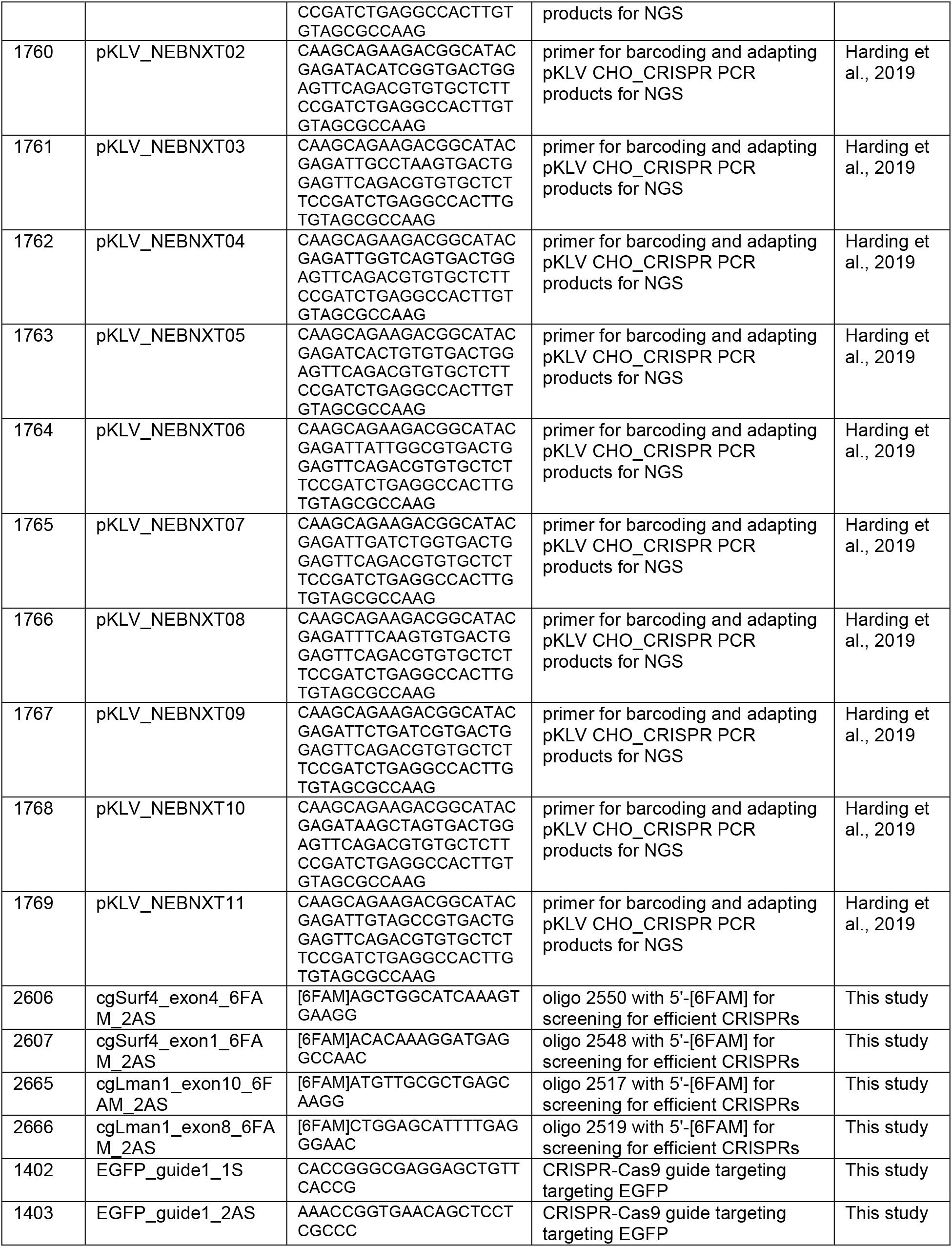
List of sgRNAs and oligonucleotides used in this study.

**Table S4:** List of antibodies, reagents and software used in this study.

| Reagent or Resource | Source | Identifier |
| --- | --- | --- |
| **Antibodies** |  |  |
| Monoclonal Mouse anti-α1AT polymer-specific 2C1 | Miranda et al., 2010 | Mab2C1 |
| Monoclonal Mouse anti-total α1AT 3C11 | Tan et al., 2015 | Mab3C11 |
| Polyclonal Rabbit anti-total α1AT | Agilent, Dako | A0012 |
| Polyclonal Rabbit anti-ERGIC-53 | Sigma | E1031 |
| Polyclonal Rabbit anti-SURF4 | Invitrogen | PA5-69676 |
| Monoclonal Mouse anti FLAG-M2 | Sigma | F1804 |
| Polyclonal Rabbit anti-cyclophilin B | Abcam | AB16045 |
| Monoclonal Mouse anti-actin | Abcam | AB3280 |
| Polyclonal Chicken anti-hamsterBiP | Avezov et al., 2013 | anti-BiP |
| Goat anti-Mouse IgG (H+L) Cross-Adsorbed Secondary Antibody, DyLight 633 | Thermo Fisher Scientific | 35513 |
| **Chemicals** |  |  |
| Doxycycline | Sigma | D9861 |
| DMEM | Sigma | D6429 |
| Tet Free Serum | Pan-Biotech | P30-3602 |
| HyClone II Serum | Thermo Fisher Scientific | SH30066.03 |
| Penicillin/Streptomycin | Sigma | P0781 |
| L-glutamine | Sigma | G7513 |
| Non-essential amino acids solution | Sigma | M7145 |
| Nutrient Mixture F12 | Sigma | N4888 |
| Lipofectamine LTX | Thermo Fisher Scientific | A12621 |
| Bafilomycin A1 | Sigma | B1793 |
| Dithiobis(succinimidyl propionate) (DSP) | Thermo Scientific Pierce | 22585 |
| Puromycin | MERCK-milipore | 540222 |
| EDTA-free Protease inhibitor Cocktail | Roche | 11873580001 |
| DMEM (-Glu/-Met/-Cys) | Gibco | 21013024 |
| Easy Tag™ Express <sup>35</sup>S Protein Labelling Mix | Perkin Elmer | NEG072007MC |
| Protein A-Sepharose | Sigma | P3391 |
| Protein G- Sepharose 4B fast flow | Sigma | P3296 |
| Anti-FLAG M2 Affinity Gel | Sigma | F3165 |
| Ni-NTA Agarose beads | Qiagen | 30210 |
| **Experimental Models: Cell lines** |  |  |
| CHO-K1 Tet-on α1AT<sup>H334D</sup> CHOP::GFP_Cas9 | This study |  |
| CHO-K1 Tet-on α1AT<sup>H334D</sup> CHOP::GFP | This study |  |
| CHO-K1 Tet-on α1AT<sup>WT</sup> CHOP::GFP | This study |  |
| CHO-K1 Tet-on α1AT<sup>H334D</sup> | Ordenez et al., 2013 |  |
| CHO-K1 Tet-on α1AT<sup>WT</sup> | Ordenez et al., 2013 |  |
| CHO-S21 | Sekine et al., 2016 |  |
| **Software and Algorithms** |  |  |
| MAGeCK | Li et al., 2014 |  |
| Metascape | Zhou et al., 2019 |  |
| MacVector |  |  |
| Photoshop and Illustrator |  |  |
| FlowJo |  |  |
| ImageJ |  |  |
| GraphPad-Prism V8 |  |  |
| Volocity V6.3 | Perkin Elmer |  |

**Table S5:**
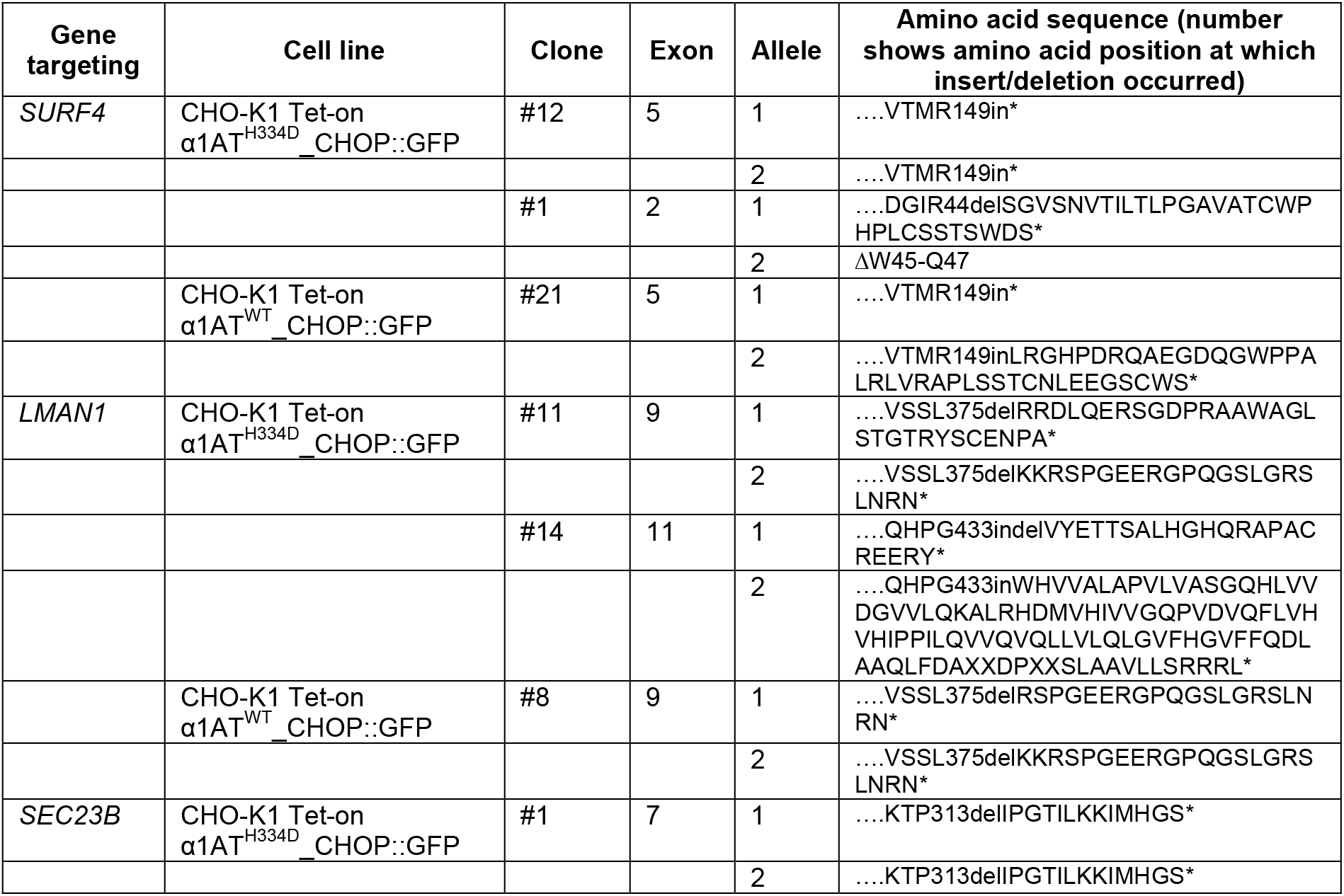
Clones generated in this study.

## References

Belden, W.J., and C. Barlowe. 2001. Role of Erv29p in collecting soluble secretory proteins into ER-derived transport vesicles. Science. 294:1528–1531.

Carrell, R.W., and D.A. Lomas. 2002. Alpha1-antitrypsin deficiency--a model for conformational diseases. The New England journal of medicine. 346:45–53.

Citterio, C., A. Vichi, G. Pacheco-Rodriguez, A.M. Aponte, J. Moss, and M. Vaughan. 2008. Unfolded protein response and cell death after depletion of brefeldin A-inhibited guanine nucleotide-exchange protein GBF1. Proceedings of the National Academy of Sciences of the United States of America. 105:2877–2882.

D’Arcangelo, J.G., K.R. Stahmer, and E.A. Miller. 2013. Vesicle-mediated export from the ER: COPII coat function and regulation. Biochimica et biophysica acta. 1833:2464–2472.

Emmer, B.T., G.G. Hesketh, E. Kotnik, V.T. Tang, P.J. Lascuna, J. Xiang, A.C. Gingras, X.W. Chen, and D. Ginsburg. 2018. The cargo receptor SURF4 promotes the efficient cellular secretion of PCSK9. eLife. 7.

Eriksson, S., J. Carlson, and R. Velez. 1986. Risk of cirrhosis and primary liver cancer in alpha 1-antitrypsin deficiency. The New England journal of medicine. 314:736–739.

Fra, A., F. Cosmi, A. Ordonez, R. Berardelli, J. Perez, N.A. Guadagno, L. Corda, S.J. Marciniak, D.A. Lomas, and E. Miranda. 2016. Polymers of Z alpha1-antitrypsin are secreted in cell models of disease. The European respiratory journal. 47:1005–1009.

Fregno, I., E. Fasana, T.J. Bergmann, A. Raimondi, M. Loi, T. Solda, C. Galli, R. D’Antuono, D. Morone, A. Danieli, P. Paganetti, E. van Anken, and M. Molinari. 2018. ER-to-lysosome-associated degradation of proteasome-resistant ATZ polymers occurs via receptor-mediated vesicular transport. The EMBO journal. 37.

Gomez-Navarro, N., and E. Miller. 2016. Protein sorting at the ER-Golgi interface. The Journal of cell biology. 215:769–778.

Gooptu, B., J.A. Dickens, and D.A. Lomas. 2014. The molecular and cellular pathology of alpha(1)-antitrypsin deficiency. Trends in molecular medicine. 20:116–127.

Gooptu, B., and D.A. Lomas. 2008. Polymers and inflammation: disease mechanisms of the serpinopathies. The Journal of experimental medicine. 205:1529–1534.

Gross, B., M. Grebe, M. Wencker, J.K. Stoller, L.M. Bjursten, and S. Janciauskiene. 2009. New Findings in PiZZ alpha1-antitrypsin deficiency-related panniculitis. Demonstration of skin polymers and high dosing requirements of intravenous augmentation therapy. Dermatology. 218:370–375.

Hauri, H.P., F. Kappeler, H. Andersson, and C. Appenzeller. 2000. ERGIC-53 and traffic in the secretory pathway. Journal of cell science. 113 (4):587–596.

Hjelm, L.N., E.L. Chin, M.R. Hegde, B.W. Coffee, and L.J. Bean. 2010. A simple method to confirm and size deletion, duplication, and insertion mutations detected by sequence analysis. The Journal of molecular diagnostics : JMD. 12:607–610.

Jensen, D., and R. Schekman. 2011. COPII-mediated vesicle formation at a glance. Journal of cell science. 124:1–4.

Kroeger, H., E. Miranda, I. MacLeod, J. Perez, D.C. Crowther, S.J. Marciniak, and D.A. Lomas. 2009. Endoplasmic reticulum-associated degradation (ERAD) and autophagy cooperate to degrade polymerogenic mutant serpins. The Journal of biological chemistry. 284:22793–22802.

Li, W., H. Xu, T. Xiao, L. Cong, M.I. Love, F. Zhang, R.A. Irizarry, J.S. Liu, M. Brown, and X.S. Liu. 2014. MAGeCK enables robust identification of essential genes from genome-scale CRISPR/Cas9 knockout screens. Genome biology. 15:554.

Lomas, D.A., D.L. Evans, S.R. Stone, W.S. Chang, and R.W. Carrell. 1993. Effect of the Z mutation on the physical and inhibitory properties of alpha 1-antitrypsin. Biochemistry. 32:500–508.

Lomas, D.A., and R. Mahadeva. 2002. Alpha1-antitrypsin polymerization and the serpinopathies: pathobiology and prospects for therapy. The Journal of clinical investigation. 110:1585–1590.

Mahadeva, R., C. Atkinson, Z. Li, S. Stewart, S. Janciauskiene, D.G. Kelley, J. Parmar, R. Pitman, S.D. Shapiro, and D.A. Lomas. 2005. Polymers of Z alpha1-antitrypsin colocalize with neutrophils in emphysematous alveoli and are chemotactic in vivo. The American journal of pathology. 166:377–386.

Miranda, E., J. Perez, U.I. Ekeowa, N. Hadzic, N. Kalsheker, B. Gooptu, B. Portmann, D. Belorgey, M. Hill, S. Chambers, J. Teckman, G.J. Alexander, S.J. Marciniak, and D.A. Lomas. 2010. A novel monoclonal antibody to characterize pathogenic polymers in liver disease associated with alpha1-antitrypsin deficiency. Hepatology. 52:1078–1088.

Mitrovic, S., H. Ben-Tekaya, E. Koegler, J. Gruenberg, and H.P. Hauri. 2008. The cargo receptors Surf4, endoplasmic reticulum-Golgi intermediate compartment (ERGIC)-53, and p25 are required to maintain the architecture of ERGIC and Golgi. Molecular biology of the cell. 19:1976–1990.

Morris, H., M.D. Morgan, A.M. Wood, S.W. Smith, U.I. Ekeowa, K. Herrmann, J.U. Holle, L. Guillevin, D.A. Lomas, J. Perez, C.D. Pusey, A.D. Salama, R. Stockley, S. Wieczorek, A.J. McKnight, A.P. Maxwell, E. Miranda, J. Williams, C.O. Savage, and L. Harper. 2011. ANCA-associated vasculitis is linked to carriage of the Z allele of alpha(1) antitrypsin and its polymers. Annals of the rheumatic diseases. 70:1851–1856.

Morrison, H.M., J.A. Kramps, D. Burnett, and R.A. Stockley. 1987. Lung lavage fluid from patients with alpha 1-proteinase inhibitor deficiency or chronic obstructive bronchitis: anti-elastase function and cell profile. Clinical science. 72:373–381.

Mulgrew, A.T., C.C. Taggart, M.W. Lawless, C.M. Greene, M.L. Brantly, S.J. O’Neill, and N.G. McElvaney. 2004. Z alpha1-antitrypsin polymerizes in the lung and acts as a neutrophil chemoattractant. Chest. 125:1952–1957.

Nyfeler, B., V. Reiterer, M.W. Wendeler, E. Stefan, B. Zhang, S.W. Michnick, and H.P. Hauri. 2008. Identification of ERGIC-53 as an intracellular transport receptor of alpha1-antitrypsin. The Journal of cell biology. 180:705–712.

Ordonez, A., E.L. Snapp, L. Tan, E. Miranda, S.J. Marciniak, and D.A. Lomas. 2013. Endoplasmic reticulum polymers impair luminal protein mobility and sensitize to cellular stress in alpha1-antitrypsin deficiency. Hepatology. 57:2049–2060.

Ran, F.A., P.D. Hsu, J. Wright, V. Agarwala, D.A. Scott, and F. Zhang. 2013. Genome engineering using the CRISPR-Cas9 system. Nature protocols. 8:2281–2308.

Reeves, J.E., and M. Fried. 1995. The surf-4 gene encodes a novel 30 kDa integral membrane protein. Molecular membrane biology. 12:201–208.

Saegusa, K., M. Sato, N. Morooka, T. Hara, and K. Sato. 2018. SFT-4/Surf4 control ER export of soluble cargo proteins and participate in ER exit site organization. The Journal of cell biology. 217:2073–2085.

Segeritz, C.P., S.T. Rashid, M.C. de Brito, M.P. Serra, A. Ordonez, C.M. Morell, J.E. Kaserman, P. Madrigal, N.R.F. Hannan, L. Gatto, L. Tan, A.A. Wilson, K. Lilley, S.J. Marciniak, B. Gooptu, D.A. Lomas, and L. Vallier. 2018. hiPSC hepatocyte model demonstrates the role of unfolded protein response and inflammatory networks in alpha1-antitrypsin deficiency. Journal of hepatology. 69:851–860.

Sekine, Y., A. Zyryanova, A. Crespillo-Casado, N. Amin-Wetzel, H.P. Harding, and D. Ron. 2016. Paradoxical Sensitivity to an Integrated Stress Response Blocking Mutation in Vanishing White Matter Cells. PloS one. 11:e0166278.

Shalem, O., N.E. Sanjana, E. Hartenian, X. Shi, D.A. Scott, T. Mikkelson, D. Heckl, B.L. Ebert, D.E. Root, J.G. Doench, and F. Zhang. 2014. Genome-scale CRISPR-Cas9 knockout screening in human cells. Science. 343:84–87.

Tan, L., J.A. Dickens, D.L. Demeo, E. Miranda, J. Perez, S.T. Rashid, J. Day, A. Ordonez, S.J. Marciniak, I. Haq, A.F. Barker, E.J. Campbell, E. Eden, N.G. McElvaney, S.I. Rennard, R.A. Sandhaus, J.M. Stocks, J.K. Stoller, C. Strange, G. Turino, F.N. Rouhani, M. Brantly, and D.A. Lomas. 2014. Circulating polymers in alpha1-antitrypsin deficiency. The European respiratory journal. 43:1501–1504.

Tan, L., J. Perez, M. Mela, E. Miranda, K.A. Burling, F.N. Rouhani, D.L. DeMeo, I. Haq, J.A. Irving, A. Ordonez, J.A. Dickens, M. Brantly, S.J. Marciniak, G.J. Alexander, B. Gooptu, and D.A. Lomas. 2015. Characterising the association of latency with alpha(1)-antitrypsin polymerisation using a novel monoclonal antibody. The international journal of biochemistry & cell biology. 58:81–91.

Walter, P., and D. Ron. 2011. The unfolded protein response: from stress pathway to homeostatic regulation. Science. 334:1081–1086.

Wu, Y., I. Whitman, E. Molmenti, K. Moore, P. Hippenmeyer, and D.H. Perlmutter. 1994. A lag in intracellular degradation of mutant alpha 1-antitrypsin correlates with the liver disease phenotype in homozygous PiZZ alpha 1-antitrypsin deficiency. Proceedings of the National Academy of Sciences of the United States of America. 91:9014–9018.

Yin, Y., M.R. Garcia, A.J. Novak, A.M. Saunders, R.S. Ank, A.S. Nam, and L.W. Fisher. 2018. Surf4 (Erv29p) binds amino-terminal tripeptide motifs of soluble cargo proteins with different affinities, enabling prioritization of their exit from the endoplasmic reticulum. PLoS biology. 16:e2005140.

Zhang, B., C. Zheng, M. Zhu, J. Tao, M.P. Vasievich, A. Baines, J. Kim, R. Schekman, R.J. Kaufman, and D. Ginsburg. 2011. Mice deficient in LMAN1 exhibit FV and FVIII deficiencies and liver accumulation of alpha1-antitrypsin. Blood. 118:3384–3391.

Zhou, Y., B. Zhou, L. Pache, M. Chang, A.H. Khodabakhshi, O. Tanaseichuk, C. Benner, and S.K. Chanda. 2019. Metascape provides a biologist-oriented resource for the analysis of systems-level datasets. Nature communications. 10:1523.

Zlatic, S.A., P.V. Ryder, G. Salazar, and V. Faundez. 2010. Isolation of labile multi-protein complexes by in vivo controlled cellular cross-linking and immuno-magnetic affinity chromatography. Journal of visualized experiments : JoVE.

